# Involvement of prelimbic cortex neurons and related circuits in the acquisition of cooperative learning by pairs of rats

**DOI:** 10.1101/2022.01.13.476162

**Authors:** A.R. Conde-Moro, F. Rocha-Almeida, E. Gebara, J.M. Delgado-García, C. Sandi, A. Gruart

**Affiliations:** Division of Neurosciences, Pablo de Olavide University, Seville-41013, Spain; Laboratory of Behavioral Genetics, Brain Mind Institute, School of Life Sciences, École Polytechnique Fédérale de Lausanne, Lausanne, Switzerland

**Author notes:** **Correspondence to:** Dr. Ana Rocío Conde-Moro, Institut Hospital del Mar d’Investigacions Mèdiques (current affiliation), Carrer del Doctor Aiguader, 88, 08003 Barcelona.

## Abstract

Social behaviors such as cooperation are crucial for mammals. A deeper knowledge of the neuronal mechanisms underlying cooperation can be beneficial for people suffering from pathologies with impaired social behavior. Our aim was to study the brain activity when two animals synchronize their behavior to obtain a mutual reinforcement. In a previous work, we showed that the activity of the prelimbic cortex (PrL) was enhanced during cooperation in rats, especially in the ones leading most cooperative trials (*leader* rats). In this study, we investigated the specific cell type/s in the PrL contributing to cooperative behaviors. To this end, we collected rats’ brains at key moments of the learning process to analyze the levels of c-FOS expression in the main cellular groups of the PrL (glutamatergic cells containing D1 and D2 receptors and interneurons). *Leader* rats showed increased c-FOS activity in cells expressing D1 receptors during cooperation. In addition, we analyzed the levels of anxiety, dominance, and locomotor behavior, finding that *leader* rats are in general less anxious and less dominant than *followers*. We also recorded local field potentials (LFPs) from the PrL, the nucleus accumbens septi (NAc), and the basolateral amygdala (BLA). Spectral analysis showed that delta activity in PrL and NAc increased when rats cooperated, while BLA activity in delta and theta bands decreased considerably during cooperation. The PrL and NAc also increased their connectivity in the high theta band during cooperation. Thus, the present work identifies the specific PrL cell types engaged in this behavior, as well as its connectivity with subcortical brain regions (BLA, NAc) during cooperation.

**Significance Statement:** Brain mechanisms underlying cooperative behaviors remain unknown. The present study identified specific neuronal types from the PrL cortex engaged in the acquisition of a cooperative task, as well as their connectivity with subcortical projection sites, such as the NAc and the BLA during cooperation. Rats leading the cooperation trials (designated *leaders*) presented an increased activation of D1-containing neurons in the PrL during cooperation. The PrL and NAc electrical activity increased when rats were cooperating, while the BLA activity increased before cooperation. The PrL and NAc showed increased functional connectivity at the moment of cooperation on the platform, whereas during the individual phase the highest connectivity was found before the animals climbed individually onto the platform.

## Introduction

In nature, many species live in groups to obtain greater benefits than by acting alone (Crawford, 1941; Eisenberg and Miller, 1987; Raihani and Bshary, 2011; Decety and Svetlova, 2012). Social behaviors, such as cooperation, are a powerful way of improving the access to resources (Crawford, 1937; Petit et al., 1992) and require a precise synchronization of animal activities (Nessler and Gilliland, 2009). For humans, social behavior is the foundational structure of our society.

Since the publication of classic social interaction studies in non-human primates (Chalmeau et al., 1997; Mendres and de Waal, 2000; Hirata and Fuwa, 2007), the number of papers focusing on prosocial behaviors has grown steadily. In the last 20 years, researchers have developed successful protocols for studying cooperative and helping behaviors in laboratory rats (Schuster and Perelberg, 2004; Rutte and Taborsky, 2007; Łopuch and Popik, 2011, Bartal et al., 2011; Hernandez-Lallement et al., 2015, Márquez et al., 2015: Sato et al., 2015). These studies have been aimed at the analysis of behavioral and cognitive strategies involved in cooperation, but information regarding the brain mechanisms underlying social behaviors and the internal drives for cooperation remains scarce.

In recent years, some researchers have recorded the brain activity —such as electrophysiology and calcium imaging—, from a variety of brain structures during social behaviors. These were mainly social approaching behaviors (Felix-Ortiz and Tye, 2014; Gunaydin et al., 2014; Lee et al., 2016; Minami et al., 2017), social competition (Zhou et al., 2017, 2018; Kingsbury et al., 2019), and helping behavior (Ben Ami Bartal et al., 2021). In a previous study, we recorded LFPs in the PrL cortex of couples of rats during the performance of a cooperation task, finding that the PrL is involved at the moment of cooperation, especially in the rats designated *leaders* —namely, the ones adjusting their behavior to that of their partner to cooperate (Conde-Moro et al., 2019).

In this work, we aimed to study which particular PrL cell types were involved in the acquisition of the cooperative task and whether the different LFP activities observed in the two groups of rats (*leaders* or *followers*) would also be confirmed at the cellular level. Four groups of rats performed the cooperation experiment, and their brains were collected at key behavioral phases to analyze c-FOS expression in the main PrL cellular subgroups. For this, we identified specific neuronal types from the PrL cortex engaged in the acquisition of a cooperative task. As a result, we observed that rats leading the cooperation trials (designated *leaders*) presented an increased activation of D1-containing neurons in the PrL during cooperation.

With the aim of determining which additional brain structures participate in the cooperation response in an additional experiment we recorded LFPs from the PrL and subcortical structures such as the NAc and the BLA, which are known to receive projections from and send projections to PrL neurons (Sessack et al., 1989; Vertes et al., 2004; Goto and Grace, 2005; Hoover and Vertes, 2007; Gabbott et al., 2005). We show here that the electrical activity of the PrL and NAc increased when rats were cooperating, while the BLA activity increased before cooperation. The PrL and NAc showed increased functional connectivity at the moment of cooperation on the platform, whereas during the individual phase the highest connectivity between these structures was found before rats climbed individually onto the platform.

## Methods

### Experimental subjects

Experiments were carried out with male Lister Hooded rats (3 months old, 250-300 g at the beginning of the experiments) provided by an authorized supplier (Charles River Laboratories, Barcelona, Spain). Upon their arrival at Pablo de Olavide Animal House (Seville, Spain), animals were housed in pairs in Plexiglas^®^ cages until the end of the experiments and were trained through all experimental phases with the same partner. Rats were randomly paired and were kept on a 12-h light/dark cycle with constant ambient temperature (21.5 ± 1 °C) and humidity (55 ± 8%). Unless otherwise indicated, animals had food and water available *ad libitum*. All the experiments were carried out following the guidelines of the European Union Council (2010/63/EU) and Spanish regulations (BOE 34/11370-421, 2013) for the use of laboratory animals in chronic experiments. Experiments were also approved by the local Ethics Committee (06/03/2018/025) of Pablo de Olavide University.

### Apparatus for social interactions

Cooperation experiments were carried out in a double Skinner box customized by our team (Fig. 1A, B) and consisting of two adjacent Skinner modules, each measuring 29.2 × 24.1 × 21 cm (MED Associates, St. Albans, VT, USA), separated by a grille partition that allowed animals to see, hear, and smell each other and to have partial physical contact (see Conde-Moro et al., 2019). Each box was equipped with a green platform (4.5 cm in height × 7 cm in width) with five infrared beams that detected when the rat was on it, a LED light, and a food dispenser where food pellets (Noyes formula P; 45 mg; Sandown Scientific, Hampton, Middlesex, UK) were delivered after the rat climbed onto the platform.

**Figure 1.**
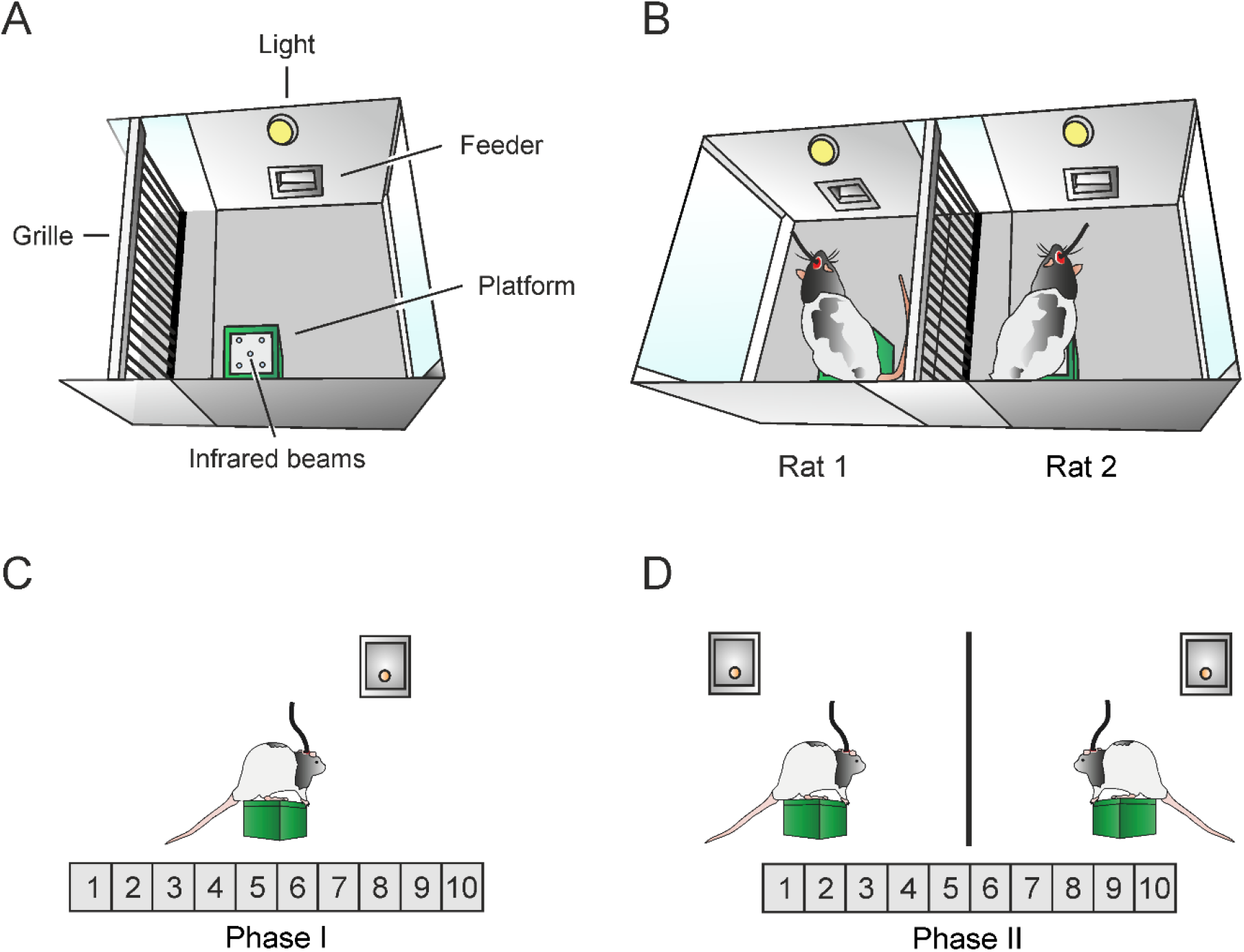
Apparatus and cooperative test. **A**, Diagram representing the details of each modified Skinner module. Each module was equipped with a food dispenser, where reinforcements were delivered, and a green platform equipped with infrared lights that detected the rat. **B**, Diagram representing the experimental setting for the cooperation experiments, where pairs of rats were placed in the double Skinner box for the two phases of the experiment. A metallic grille that allowed partial physical contact separated the two Skinner modules. **C**, Cooperative test. In phase I, animals were trained to individually climb onto a platform and stay on it for > 500 ms to get a food pellet in a fixed ratio schedule (1:1). **D,** In phase II, animals were trained to climb onto the platforms and to stay on them simultaneously for > 500 ms to get a food pellet for each of them. Training sessions lasted for 20 min. The number of experimental sessions is indicated in gray. below the diagrams.

Before experiments, rats were handled daily for 10 days and food-deprived to 85% of their free-feeding weight. Once the goal weight was reached, rats were habituated to the test room for two 10-minute free-exploration sessions in an empty Plexiglas^®^ box, different from the experimental box. Between use for each pair of animals, apparatus and cages were cleaned with 5% ethanol and dried with paper.

### Cooperation experiment

During phase I, pairs of animals were placed in the adjacent Skinner boxes for 20 min and were free to explore the cage (Fig. 1C). Each time an animal stayed on the platform for > 500 ms, a pellet of food was delivered to the feeder, following a fixed-ratio schedule (FR 1:1) until the established criterion was reached —that is, to climb onto the platform ≥ 60 times/session for two consecutive days. A LED light located above the feeder indicated when a pellet of food was delivered.

For phase II (simultaneous cooperative task), within the same set-up, animals had to climb and stay on their platforms simultaneously for > 500 ms to obtain a mutual reward (Fig. 1D). If either of the animals climbed onto the platform and went to the feeder on its own, the trial was considered wrong (for acting individually) and there would be no reward for either of them. Animals were trained daily until reaching the established criterion —that is, to climb simultaneously onto the platform to get the mutual reward ≥ 40 times/session for at least two consecutive days. Conditioning programs, LFP recordings, platform climbs, and delivered reinforcements were monitored and recorded. All room lights, except for dim light, were switched off before each experimental session to improve the rats’ comfort.

### Experimental design

Two cohorts of rats participated in this study. The objective with the first cohort, consisting of 40 animals (38 at the end of the experiment), was to study the effects of the cooperation experiment and the different roles developed by each rat (*leader* or *follower*) in the main cell types of the PrL cortex, using immuno-colocalization analysis. The objective with the second cohort, consisting of 18 animals (10 at the end of the experiment with good recordings), was to study the effect of cooperation not only in the PrL but also in subcortical projection areas (NAc and BLA), using *in vivo* recording and analysis of the LFPs collected in the aforementioned areas.

### Surgery

To prepare animals for the *in vivo* electrophysiological experiment, rats were anesthetized with 1-2.5% isoflurane delivered by a rat anesthesia mask (David Kopf Instruments, Tujunga, CA, USA). Isoflurane was supplied from a calibrated Fluotec 5 (Fluotec-Olmeda, Tewksbury, MA, USA) vaporizer, at a flow rate of 1–3 L/min oxygen (AstraZeneca, Madrid, Spain).

For LFP recordings, and following the Paxinos and Watson atlas (2007), animals were chronically implanted with two sets of recording electrodes aimed at the right PrL cortex (3.24 mm anterior, 0.5 mm lateral to bregma, and 2.5 mm from brain surface) and the right NAc Core (2.0 mm anterior and 1.5 mm lateral to bregma, and 6.5 mm from brain surface) and one set of recording electrodes aimed at the BLA (2.28 mm posterior and 5 mm lateral to bregma, and 7.5 mm from brain surface).

All electrodes were handmade from 50 μm, Teflon-coated, tungsten wire (Advent Research, Eynsham, UK). Each electrode set consisted of two tungsten wires with a separation between tips of ≈ 0.3 mm. The Teflon coating was removed from the first 200 μm of each cable tip for better wire surface exposure. A bare silver wire was affixed to the bone as ground. All the implanted wires were soldered to sockets (RS Amidata) that were fixed to the skull with six small bone anchor screws (Stoelting Co., Woodale, IL, USA) and dental cement.

### Perfusion and histology

#### Nissl staining

At the end of the experiments, rats were deeply re-anesthetized with a mixture of ketamine (100 mg/kg) and medetomidine (0.1 mg/kg) and perfused transcardially with saline and 4% paraformaldehyde in phosphate-buffered saline (PBS, 0.1M, pH 7.4). Brains were cryoprotected with 30% sucrose in PBS for a few days, after which 50 μm coronal sections were obtained with a sliding freezing microtome (Leica SM2000R, Nussloch, Germany). Selected sections that included the implanted areas were mounted on gelatinized glass slides and stained using the Nissl technique with 0.1% toluidine blue to reveal the final location of recording electrodes in the PrL cortex, NAc, and BLA (Fig. 1-1).

#### Brain collection and preparation for immunofluorescence

To collect the brain tissue at key moments of the cooperation acquisition, three groups started the cooperation experiment and rats from each group were sacrificed at a different point of the experiment (Fig. 2A).

**Figure 2.**
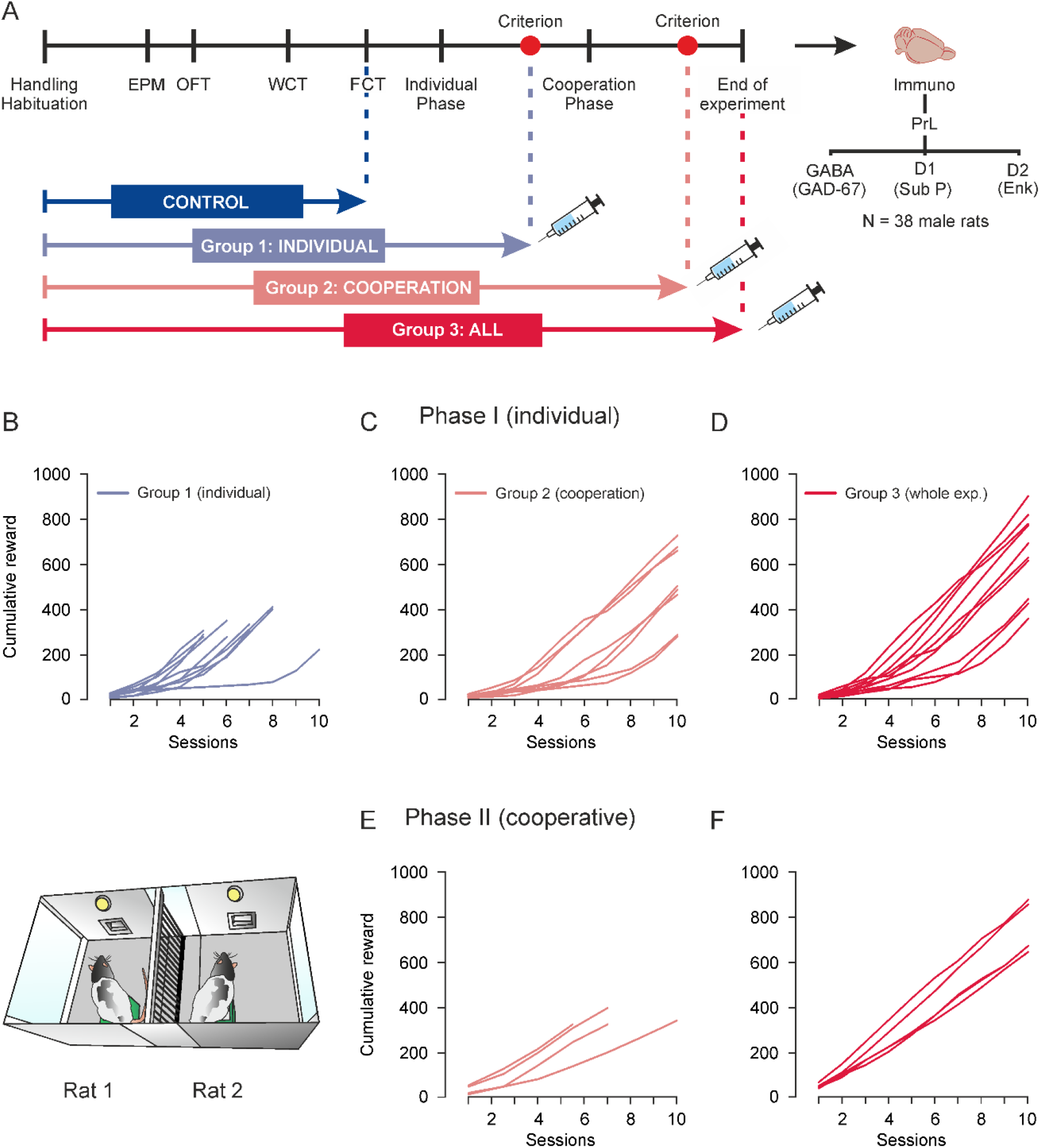
Immunofluorescence procedure and cooperation performance. **A, Immunofluorescence procedure.** Animals (n = 40) were randomly allocated into 4 groups of 10 rats: control group (blue), group 1 (purple), group 2 (pink), and group 3 (dark red). All animals underwent a battery of behavioral tests before the cooperation experiment: open field test (OFT), elevated plus maze (EPM), water competition test (WCT), and food competition test (FCT). After that, groups 1, 2, and 3 started the cooperation experiment. The brains of each group were collected after reaching the established key moments (indicated by syringe icons in the image): brains of group 1 were collected after reaching the criterion for the individual phase (i.e., ≥ 60 pellets for 2 consecutive days), brains in group 2 after reaching the criterion for the cooperative phase (i.e., ≥ 40 pellets for 2 consecutive days), and brains of group 3 at the end of the experiment. Animals in the control group remained in their home cages during the cooperation experiment and were handled and weighed daily as the remaining animals. After the cooperation experiment, all the brains were cryoprotected, and coronal sections of the PrL cortex were labeled for c-FOS, DAPI, and selected antibodies against GABAergic and dopaminergic neurons (D1- and D2-containing cells). **B-F**, Cumulative records showing the acquisition curves for individual and cooperative phases. 38 male rats were randomly allocated into 4 groups (group 1 (individual phase, represented in purple), group 2 (cooperative phase, represented in pink), group 3 (whole experiment, represented in red), and the control group. All groups except the control performed the cooperation experiment. **B**, Group 1 learned to individually climb and sit on the platform to get a pellet of food, showing a steep slope between the third and the last day of training: average ± SD slope for group 1, 76.801 ± 22.46. The cumulative line stops at the day that rats reached the criterion and were perfused and euthanized. **C**, **D**, Groups 2 and 3 performed the individual phase completely and all the animals reached the criterion, showing a steep slope from day 3 to 10 (group 2, slope = 77.03 ± 26.01; and group 3, slope = 94.24 ± 27.45). **E**, **F**, After learning the individual task, groups 2 and 3 were placed again in the double Skinner box for phase II (cooperative). This time, they had to coordinate their behavior (climbing and staying together on the platform for > 500 ms) to get a pellet of food. All rats learned the task successfully, showing a steep slope from session 3 to their last session (group 2, slope = 87.82 ± 17.31; group 3, slope = 82.45 ± 14.04).

Group 1 was the individual group, which performed the protocol until reaching the criterion for the individual phase (i.e., getting ≥ 60 pellets of food for two consecutive days). Group 2 was the cooperation group, which performed all phases of the protocol until they reached the criterion for the cooperation phase (i.e., ≥ 40 pellets for two consecutive days), and Group 3 completed the whole experimental protocol (i.e., they completed the 10 sessions of the cooperation phase even after reaching criterion). The control group, which underwent the same protocols as the remaining rats but did not perform the cooperation experiments, were sacrificed in a scattered way, distributed along with the perfusion of the other groups (i.e., 3 rats perfused along with group 1, 3 rats with group 2, and 4 rats with group 3, always counterbalancing the order with those from experimental groups). Rats from the control group were handled 2 min per day on the same days that the other animals were performing experiments and were perfused 100 min after the first opening of the cage for handling. The order in which each pair started the experiments every day was counterbalanced.

As explained before, rats performed the cooperation experiment until they reached the criterion for their group. The day after reaching criterion, animals performed a reminder session of 15 min, and 85 min later were anesthetized (IP) with pentobarbital (150 mg/kg) and sacrificed by transcardial perfusion using 0.9% saline solution.

After perfusion, rats’ brains were carefully removed and kept in PFA 4% at 4 °C for one day and put into a sucrose solution (30%) at 4 °C for 3 days or until brains sank in the solution. When brains were ready, they were frozen with liquid nitrogen and stored at −80 °C. Afterward, brain areas of interest were serially cut at the cryostat (Leica, CM3050 S) in coronal sections (30 μm thick) and kept in a cryoprotectant solution (30% glycerol, 30% ethylene glycol, and 0.2M phosphate buffer) at −20 °C.

### Immunofluorescence

Free-floating sections were labeled for c-FOS, DAPI, and one of the antibodies against the target cell types of the PrL cortex. Sections were washed three times in 0.1M phosphate buffer and blocked with 0.1M phosphate buffer containing 10% Triton (VWR, M143-1L) and 5% normal donkey serum (Merck Millipore S30-100mL). After that, sections were incubated for 40 h at 4 °C with rabbit anti-c-FOS (Synaptic Systems, 226003) or goat anti-c-FOS (Santa Cruz, sc-52-G) and one of the following primary antibodies: mouse anti-GAD67 for staining GABAergic cells (Merck & Co., MAB5406), goat anti-Substance P for staining D1-containing cells (Santa Cruz, sc-9758), and rabbit anti-Met Enkephalin for staining D2-containing cells, (Abcam, ab22620). The sections were then incubated for 2 h in one of the secondary antibodies: donkey anti-mouse Alexa 647 (Thermo Fisher Scientific, A-31571) for GAD67, donkey anti-goat Alexa 488 (Thermo Fisher Scientific, A-11055) for Substance P, donkey anti-rabbit Alexa 568 (Thermo Fisher Scientific, A-10042) for Met-Enkephalin, and donkey anti-rabbit Alexa 568 (Thermo Fisher Scientific, A-10042) or donkey anti-goat Alexa 488 (Thermo Fisher Scientific, A-11055) for c-FOS. After washing in 0.1M phosphate buffer, the sections were incubated for 10 min in DAPI (Sigma Aldrich), rinsed, and mounted with Fluoromount-G (Southern Biotech). Images were taken with a confocal microscope (Zeiss, LSM700) using a 20x objective and were captured at the same coordinates for each animal (see Hollis et al., 2015).

For immunofluorescence quantification, we used the software FIJI. We calculated each channel background by measuring four random background areas and calculating the mean, which was subtracted from each channel. Cells were delineated with a Huang threshold to label those cells stained with DAPI within 50-200 pixels. Once we had this number of cells, we counted how many of them were also labeled with c-FOS and the antibody of interest, and converted it to a percentage.

### Anxiety, social dominance, and locomotor activity

To find out behavioral traits that could predict which rats from the groups that did not complete the cooperative phase of the experiment (control group and group 1) would develop a *leader* or *follower* role in the future (denominated here as *predicted leaders* and *predicted followers*), all animals of the immunofluorescence experiment, including the control group, underwent a battery of tests to measure their basal levels of anxiety, locomotor behavior, and social dominance before the cooperation experiment.

#### Open field test (OFT)

The OFT is used to assess anxiety-like and locomotor behaviors. It consists of a walled, square open arena (42 × 42 × 34.2 cm). The animal is placed near the wall and can explore the arena for 10 min. For analysis, the arena is divided into 3 parts: a center zone unprotected and more anxiogenic for the animals, an intermediate zone, and an outer zone, the closest to the walls and least anxiogenic for the animals. After 10 min, a novel object is placed in the center of the open field arena. The parameters analyzed for this study were the total distance traveled and the percentage of time spent in each zone. Between use for each animal, the arena was cleaned with 5% ethanol and dried with paper.

#### Elevated plus maze (EPM)

The EPM is used to assess anxiety-like behavior. This metallic apparatus consists of an elevated platform (71 cm above the ground) with arms shaped like a plus sign. The two closed arms (49 × 10 × 40 cm) are protected by walls, and the two opposing open arms (49 × 10 cm) are unprotected, which is more anxiogenic for the animals. The closed and open arms are connected by a central square area (10 × 10 cm). At the beginning of the test, animals were individually placed in the central area of the maze, facing a closed arm, and let explore the apparatus for 5 min. After every trial, the maze was cleaned with 5% ethanol and dried with paper. The parameters analyzed for this experiment were the time spent in the open and closed arms and the center zone.

#### Water and Food competition tests (WCT, FCT)

To measure social dominance, we used the water and food competition tests, in which animals compete for resources after a time of deprivation (Cordero and Sandi, 2007). The water competition test (WCT) is used to assess social dominance between rats. The experiments took place in the housing cages where rats lived in cohabitation for at least 1 week before the test. Animals were water-deprived for 6 hours before the test and marked on the back for identification. A single bottle of water was placed in an accessible part of the cage at the beginning of the test, and the rats’ behavior recorded for 10 min after the bottle is introduced.

For the food competition test (FCT), which also took place in the home cages, the protocol was similar to the water competition test, but instead of water, we placed 10 pellets of palatable food (Noyes formula P; 45 mg; Sandown Scientific, Hampton, Middlesex, UK) in the middle of the cage. Rats were food-deprived for 12 hours before the test.

The WCT and FCT sessions were also recorded and social behaviors such as aggression or displacements from the feeder or water bottle were computed. Social dominance was determined by the summation of the total duration of water consumption and the number of food pellets eaten by each rat in the pair.

### Statistical analyses

Data for quantification of animal performance in the Skinner boxes were selected and analyzed offline using Spike 2 software (Cambridge Electronics Design) and statistically analyzed afterwards using Sigma Plot 11. LFP power spectra, spectrograms, coherence spectra, and coherograms from different groups and conditions were statistically compared using Chronux customized scripts (Mitra and Bokil, 2008; Bokil et al., 2010) to obtain the jackknife estimates of the variance and of Z-statistics (for details refer to Bokil et al., 2007).

For multivariate statistics assessments, both parametric [One-way ANOVAs, with and without repeated measures (RM)] and non-parametric [One-way ANOVA tests on ranks, with and without RM (Kruskal-Wallis ANOVA)] methods were used to evaluate the statistical significance of differences between groups, followed by the appropriate test (Holm-Sidak, Tukey, or Student-Newman-Keuls, in this order of priority) for all the pairwise multiple-comparison analyses. When the normality (Shapiro-Wilk test) and equal variance of the errors (Levene Median test) assumptions were satisfied, the significance (p-value) and the statistic F were reported.

When the normality assumption was not verified, the significance (p-value) of the Chi-square (χ2) was calculated using the ranks of the data rather than their numeric values. Also, the H-statistic (One-way ANOVA on ranks between two groups) and the sample size of data were used to estimate the corresponding effect size index. Finally, Z-tests were used to compare categorical variables.

Unless otherwise indicated, data are represented by the mean ± SEM. For all the statistical tests, the significance level (p-value) is indicated. The asterisks in the graphs denote statistical significance: p-value < 0.05 (*), < 0.01 (**), or < 0.001 (***).

### Data collection, analysis and representations

#### Behavioral data collection

Data and videos of animal performance in the Skinner boxes were acquired online using the Spike 2 software (Cambridge Electronics Design). One-volt rectangular pulses corresponding to platform climbs, and pellet delivery were stored digitally on a computer for posterior analysis in conjunction with the LFP data. Locomotor activities, and EPM and OFT sessions were video recorded and analyzed with the help of Ethovision 11.0 XT software and camera by Noldus Information Technology and scored with The Observer 11.5 XT software, also by Noldus. WCT and FCT sessions were recorded with a handy camera (Sony HDR-SR12E, Tokyo, Japan) and analyzed also with The Observer XT 11.5.

#### In vivo LFP recordings

LFP activities were recorded with Grass P511 differential amplifiers with a bandwidth of 0.1 Hz – 3 kHz (Grass-Telefactor, West Warwick, RI, USA) through a high-impedance probe (2 × 1012 Ω, 10 pF) and stored digitally on a computer through analog-to-digital converters (CED 1401 Plus; Cambridge Electronics Design). LFPs were sampled at 5 kHz with an amplitude resolution of 16 bits. For analysis, we selected 2-second LFP epochs, collected from the two experimental phases (individual and cooperative), and two experimental conditions (BEFORE- and ON-platform).

#### LFP spectral decomposition and representation

The computational tools used for neurophysiological signal processing and analysis were customized MATLAB scripts (version 9.4, R2018a. The MathWorks, Natick, MA, USA) based on Chronux, (version 2.12, 2016. Website: http://chronux.org/) by Mitra and Bokil (2008). Chronux is a spectral analysis toolbox for MATLAB specialized in the processing of brain signals, which has been validated and widely used for experimental procedures, including LFPs.

Analyses in the frequency domain were carried out according to the following frequency bands: delta (3–6 Hz), low theta (6–9 Hz), high theta (9–12 Hz), beta (12–32 Hz), and gamma (32–100 Hz). A high-pass filter was applied to remove low-frequency (0–2 Hz) movement artifacts. A band-pass filter (0-200 Hz) was also applied to remove artifacts produced by the animal’s chewing.

Analyses in the frequency and time-frequency domains and coherence described in this section were computed following the multi-taper spectral estimation method developed by Thomson (1982) and implemented in the Chronux toolbox. The spectral power estimates the magnitude of Fourier transforms in the frequency domain, highlighting the frequencies with higher energy (power) within the signals. Spectral power was computed for the epochs selected for each phase and condition. Multi-taper spectrograms were also computed for all phases and conditions to represent Fourier coefficients in the time-frequency domains, which allowed us to inspect the moment at which significant changes in power took place. Spectrograms were computed using a moving time window (T) with a length of 500 ms (shifted in 10-ms increments) and bandwidth (W) of 6 Hz, resulting in a bandwidth product of 3 and K = 5 tapers. These parameters verify the number of selected tapers (i.e., K = 2 × T × W – 1 taper or windowing functions). The multi-taper estimates of the spectrum with NT trials and K tapers were based on computing NT × K Fourier transforms that determined an appropriate number of degrees of freedom; dof = 2 × NT × K for all the computations (see details in Conde-Moro et al., 2019).

To study the functional connectivity between the LFPs recorded from different brain structures, we computed the LFP-LFP phase coherence and coherograms, revealing the levels of oscillatory synchrony between electrodes from different structures at certain frequencies.

## Results

### Immunofluorescence experiments

#### All groups successfully completed the cooperation experiment

As described in Methods, 28 rats —group 1, group 2 and group 3— were successfully trained in pairs in adjacent Skinner boxes for the individual phase (phase I) and 18 rats —groups 2 and 3— were trained for the cooperation phase (phase II).

During phase I, rats were trained to climb independently —regardless of their partner’s behavior— onto the platform to obtain a food pellet at a fixed 1:1 ratio. As illustrated in the cumulative records (Fig. 2B-D), all rats improved their performance across sessions. For the majority of rats, the response onset started at session 3 and continued increasing from that session on, showing a steep slope between the third and the last training days: average ± SD slope for group 1, 76.801 ± 22.46; group 2, 77.03 ± 26.01; and group 3, 94.24 ± 27.45. Although rat #12 (pair 6) of group 1 (slope = 29.83) and rats #27 and #28 (pair 14) of group 2 (slopes 41.5 and 41.33 respectively) maintained a flatter curve until session 7, they increased their response in the last two or three sessions. All rats reached criterion from sessions 5 to 10. Rats in group 1 (individual) were anesthetized and sacrificed the day they reached criterion (see Methods for details).

During phase II, the remaining rats (groups 2 and 3) were trained to climb onto their respective platforms and stay on them simultaneously for at least 500 ms to mutually get a reward (a food pellet for each rat). As shown in the cumulative reward graphs (Fig. 2E, F), all pairs of rats learned to climb simultaneously onto the platform to mutually obtain a reward and reached the selected criterion between sessions 4 and 10. The response onset for most pairs of animals was also observed from session 3, and the slopes calculated revealed a steep increase in the number of responses across days (group 2, 87.82 ± 17.31; group 3, 82.45 ± 14.04).

As in our previous study (Conde-Moro et al., 2019), the 18 rats that performed the cooperation phase —groups 2 and 3— also adopted different strategies. For each pair, the rat that climbed onto the platform significantly more times in first place, and thus initiated more cooperation trials, was classified as *leader* (Fig. 3A, One-way ANOVA, F = 14.40, p = 0.002), while the partner was classified as *follower*. Although in that work the number of platform climbs was similar for *leaders* and *followers*, this time *leader* rats did significantly more platform climbs (Fig. 3B, One-way ANOVA, F = 18.0, p < 0.001). In addition, *leader* rats did fewer wrong trials than the *followers*, but the difference was not statistically significant (Fig. 3C, One-way ANOVA, F = 1.07, p = 0.318).

**Figure 3.**
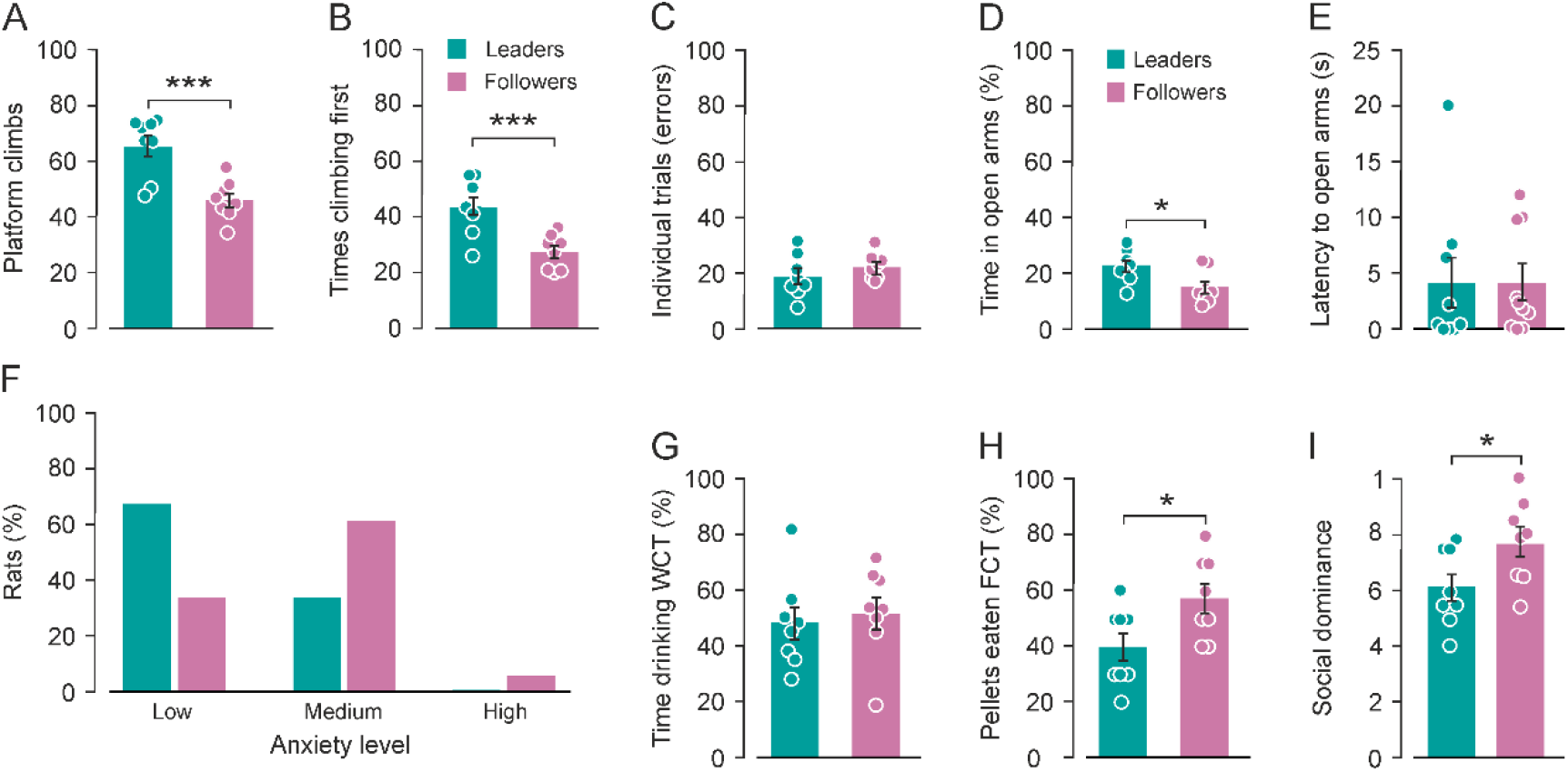
*Leader/follower*, anxiety and social dominance. **A-C**, *Leader/follower* strategy to cooperate. **A,** Average number of times that *leader* and *follower* rats climbed in first place onto the platform during the cooperative phase. *Leader* rats climbed significantly more times in first place (One-way ANOVA, F = 14.40, p = 0.002), initiating the cooperation trials more times. **B**, Average number of platform climbs during Phase II (cooperation) of *leaders* (in green) and *follower* rats (in purple). *Leader* rats climbed significantly more times onto the platform than *follower* rats (One-way ANOVA, F = 18.00, p < 0.001). **C**, Average number of wrong trials for *leader* and *follower* rats. Although there is a tendency for *leader* rats to make fewer wrong responses, there were no significant differences between them (One-way ANOVA, F = 1.073, p = 0.318). **D-E**, Anxiety levels (groups 2 and 3). **D**, Average time (percentage) spent in the open arms of the elevated plus maze (EPM). *Leader* rats spent significantly more time than *followers* in open arms (One-way ANOVA, F = 6.71, p = 0.02), which indicates lower levels of anxiety. **E**, Latency to enter any of the open arms (in seconds) for *leader* and *follower* rats. No significant differences were found between *leaders* and *followers* in this case (One-way ANOVA on ranks, H = 0.24, p = 0.62). **F**, percentage of *leader* and *follower* rats that fell into the three categories of anxiety established: low-anxious (LA, more than 20% of the time spent in the open arms of the EPM), medium-anxious (MA, between 5% and 20% of the time in open arms), and high-anxious (HA, less than 5% of the time in open arms). Most *leader* rats were classified as low-anxious (66%), while most *follower* rats were classified as intermediate-anxious (61%). **G-H,** Social dominance (groups 2 and 3). **G**, Percentage of time drinking water during the water competition test. *Leaders* and *followers* spent a similar percentage of time drinking water (One-way ANOVA, F = 0.18, p = 0.67). **H**, Percentage of food pellets eaten during the food competition test. *Follower* rats ate a significantly higher percentage of food pellets than *leaders* (One-way ANOVA, F = 8.69, p = 0.01). **I**, Social dominance index, based on the percentage of time drinking water, the percentage of food pellets eaten, and the number of aggressions and successful displacements from the water bottle or feeder. *Follower* rats showed a significantly higher level of social dominance (One-way ANOVA, F = 5.29, p = 0.03) than *leader* rats during these tests.

#### Leader rats were less anxious than follower rats

Before the cooperation experiment, the four groups of rats (n = 38) underwent a battery of validated behavioral tests in an attempt to assess behavioral traits, such as anxiety (EPM test), locomotor activity (OFT), and social dominance (WCT and FCT).

Although *leader* rats, in general, spent more time in the center area of the arena than *followers*, they presented a high variability, and no significant differences between *leader*s and *followers* were found in the percentage of time spent in the center —and more anxiogenic— area of the OFT (Fig. 2-1A, One-way ANOVA, F = 2.58, p = 0.12). No significant differences were found either in the total distance traveled by *leaders* and *followers* during this test (Fig. 2-1B, One-way ANOVA, F = 1.95, p = 1.81). After 10 min in the open field arena, a novel object was inserted in the center for 5 min. Again, there were no significant differences between the percentage of time that *leader* and *follower* rats spent in the object zone (Fig. 2-1C, One-way ANOVA, F = 1.10, p = 0.30).

However, *leader* rats spent significantly more time in the open arms of the elevated plus-maze (Fig. 3D, One-way ANOVA, F = 6.71, p = 0.02), showing lower levels of anxiety. No significant differences were found in the latency to enter the open arms (Fig. 3E, One-way ANOVA on ranks, H = 0.24, p = 0.62). According to the scale validated by Sandi et al. (2008), rats can be classified by three different levels of anxiety based on the percentage of time spent in the open arm of the EPM: low-anxious (LA, more than 20% of the time in the open arms), intermediate-anxious (IA, between 5% and 20% in the open arms), and high-anxious (HA, less than 5% in the open arms). Most *leader* rats in our study were classified as low-anxious (Fig. 3F, LA = 66%, IA = 33%); no rat in the *leader* group was classified as high-anxious. In the *follower* group, most of the rats were classified as intermediate-anxious (Fig. 3F, LA = 33%, IA = 61%, HA = 5%). These results indicate that *leader* rats were in general less anxious than *follower* rats.

#### Follower rats showed higher social dominance than leader rats

For this experiment, we used WCTs and FCTs, in which animals compete for resources (water or food) after a time of deprivation. To measure social dominance, we calculated the percentage of time that rats spent drinking water during the water competition test and the percentage of food intake during the food competition test. *Leader* and *follower* rats spent a similar percentage of time drinking water in the water competition test (Fig. 3G, One-way ANOVA, F = 0.18, p = 0.67), while *follower* rats ate a significantly higher percentage of food pellets during the water competition test (Fig. 3H, One-way ANOVA, F = 8.69, p = 0.01).

To have a deeper insight of the hierarchical dynamics taking place, we summed the number of successful displacements and aggressions to the social dominance index based on the work of Costa, et al., (2021). We found that *follower* rats showed a significantly higher level of social dominance (Fig. 3I, One-way ANOVA, F = 5.29, p = 0.03) than *leader* rats during these tests.

#### c-FOS expression was higher in PrL D1-containing cells of leader rats during cooperation

To determine the level of activation of putative neuronal groups involved in cooperation behaviors (D1- and D2-containing cells and GABAergic cells), 3 groups of rats participated in the cooperation experiment; they were sacrificed at key moments and their brain tissue was collected. Group 1 participated in the protocol until reaching criterion for the individual phase, group 2 participated in the protocol until reaching criterion for the cooperation phase, and group 3 completed the whole protocol (10 sessions of individual training and 10 of cooperative training). A control group participated in the previous behavioral tests (i.e., open field, elevated plus) but not in the cooperation experiment (see Methods section).

Analysis of the percentage of c-FOS expression in PrL cortex dopaminergic cells revealed a significantly higher activation of D1-containing cells of *leader* rats from group 2 during the cooperation phase (Fig. 4A, B, One-way ANOVA, F = 16.95, p = 0.007), while non-significant differences were found between *leaders* and *followers* from group 3 (Fig. 4C, D, One-way ANOVA, F = 5.33, p = 0.08), although the activation level of D1-containing cells was higher for *leader* rats. The levels of c-FOS expression in D2-containing cells were lower than those in D1-containing cells.

**Figure 4.**
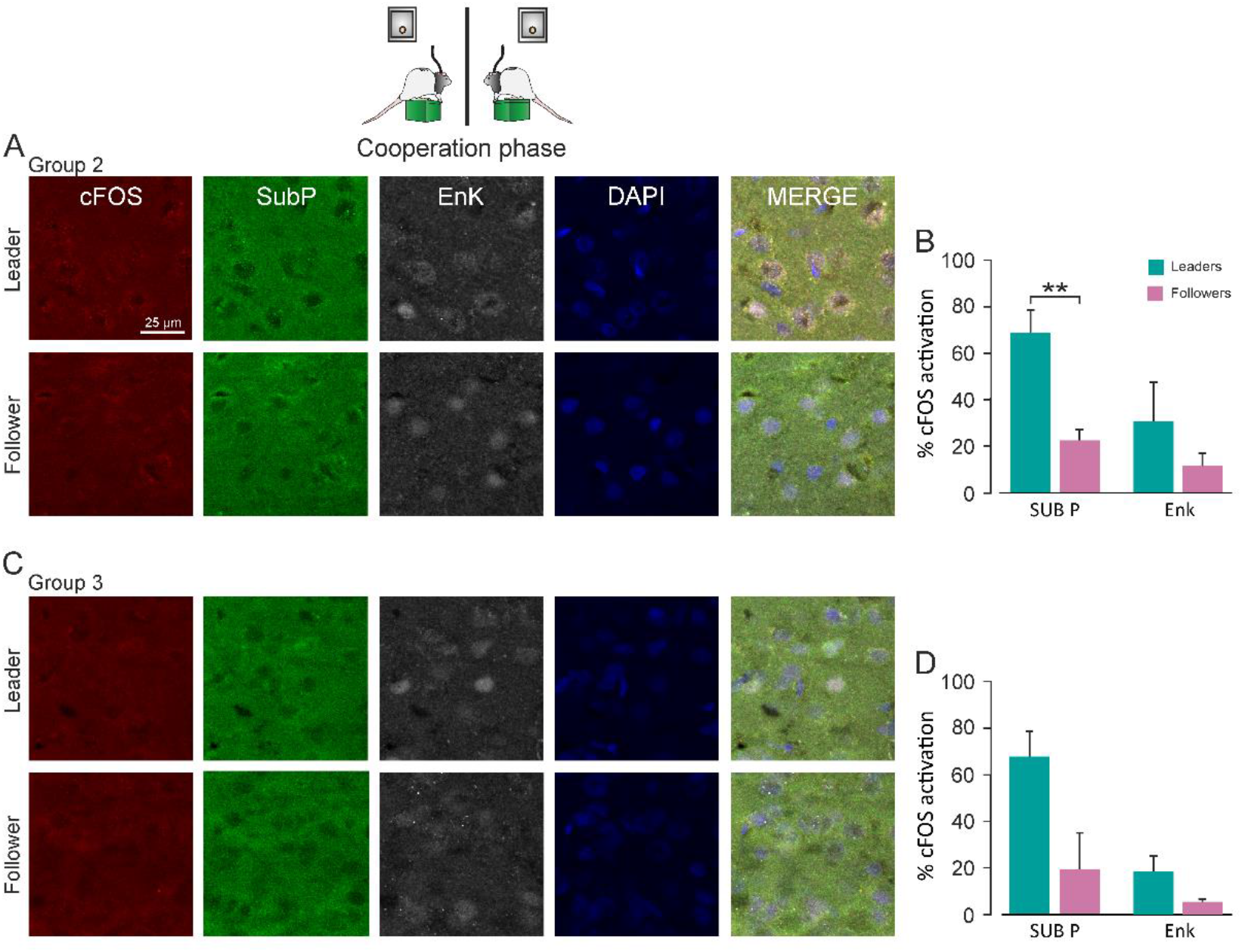
c-FOS expression in PrL D1- and D2-containing cells of cooperation groups (groups 2 and 3). **A**, **C**, Coronal brain sections from animals in groups 2 and 3 labeled for c-FOS (red channel), DAPI (blue channel), and the selected antibodies against the main cell types of the PrL. Goat anti-Substance P for staining D1-containing cells (green channel) and rabbit anti-Enkephalin for staining D2-containing cells (gray channel). The photomicrographs corresponding to Group 2, in which rats were trained to cooperate until reaching the criterion for the cooperation phase, are shown in **A,** and the photomicrographs in **C** correspond to group 3, which performed the whole cooperative phase (10 sessions). **B**, **D**, Percentage of c-FOS activation in each group. **B**, D1-containing cells were significantly more active in *leaders* than in *followers* during the cooperation phase (One-way ANOVA, F = 16.95, p = 0.007). The activation of D2-containing cells was lower than that of D1 cells, and the highest activation was also observed in the *leader* rats from group 2. However, the difference between *leaders* and *followers* was not significant (One-way ANOVA, F = 1.63, p = 0.24). *Leader* rats from group 3 (**C**), which completed the 10 days of the task, also showed higher activation for D1 and D2 cells, but the difference was not significant (One-way ANOVA, F = 1.44, p = 0.29).

The highest levels of activation of D2-containing cells were also observed for *leader* rats during the cooperation phase (Fig. 4A, B), although the difference between *leader* and *follower* rats was not significant (Fig. 4B, One-way ANOVA, F = 1.63, p = 0.24; D, One-way ANOVA, F = 1.44, p = 0.29).

During the individual phase, and contrarily to what happened during cooperation, the rats predicted to be *followers* (*predicted followers*) presented higher percentages of c-FOS expression than the ones predicted to be *leaders* (*predicted leaders*), but this difference was non-significant (Fig. 4-1C, D, One-way ANOVA, F = 0.68, p = 0.42). The same happened in the control group (Fig. 4-1A, B, One-way ANOVA on ranks, H = 0.02, p = 0.93). Non-significant differences were found between the level of activation of D2 cells for *predicted leaders* and *predicted followers* during the individual phase (Fig. 4-1D, One-way ANOVA on ranks, H = 2.41, p = 0.11) and in the control group (Fig. 4-1B, One-way ANOVA, F = 1.85, p = 0.20).

Regarding the percentage of c-FOS activation in GABAergic cells during the cooperative phase, although *leader* rats presented higher levels of activation than *followers*, non-significant differences were found in the levels of c-FOS expression for GABAergic cells between *leaders* and *followers* of groups 2 (Fig. 4-2A, B, One-way ANOVA, F =2.50, p = 0.12) and 3 (Fig. 4-2C, D, One-way ANOVA, F = 1.62, p = 0.22).

In GABAergic cells, the levels of c-FOS expression were higher in *predicted follower* rats during the individual phase and in the control group, but, again, the differences between *predicted leaders* and *predicted followers* were not significant (Fig. 4-3D, One-way ANOVA, F = 2.15, p = 0.16; B, One-way ANOVA, F = 0.49, p = 0.49).

### Electrophysiological experiments

#### All groups successfully completed the cooperation experiment

As described in Methods, during the cooperation experiment we analyzed LFPs recorded from three different structures of the brain: the PrL cortex, the NAc, and the BLA. A total of 10 rats were successfully trained pair-wise in the adjacent Skinner boxes for the individual phase (phase I) and the cooperation phase (phase II).

During phase I, rats were trained to climb independently —regardless of their partner’s behavior— onto the platform to obtain a pellet of food at a fixed 1:1 ratio. As illustrated in the acquisition curve (Fig. 5A), rats improved their performance across sessions. Compared with session 1, the average number of correct responses increased significantly from session 4 on (RM-One-way ANOVA, multiple comparisons: S04, p < 0.05; S05, S06, S07, p < 0.01), showing an even higher increase in the last three sessions (S08, S09, S10, p < 0.001).

**Figure 5.**
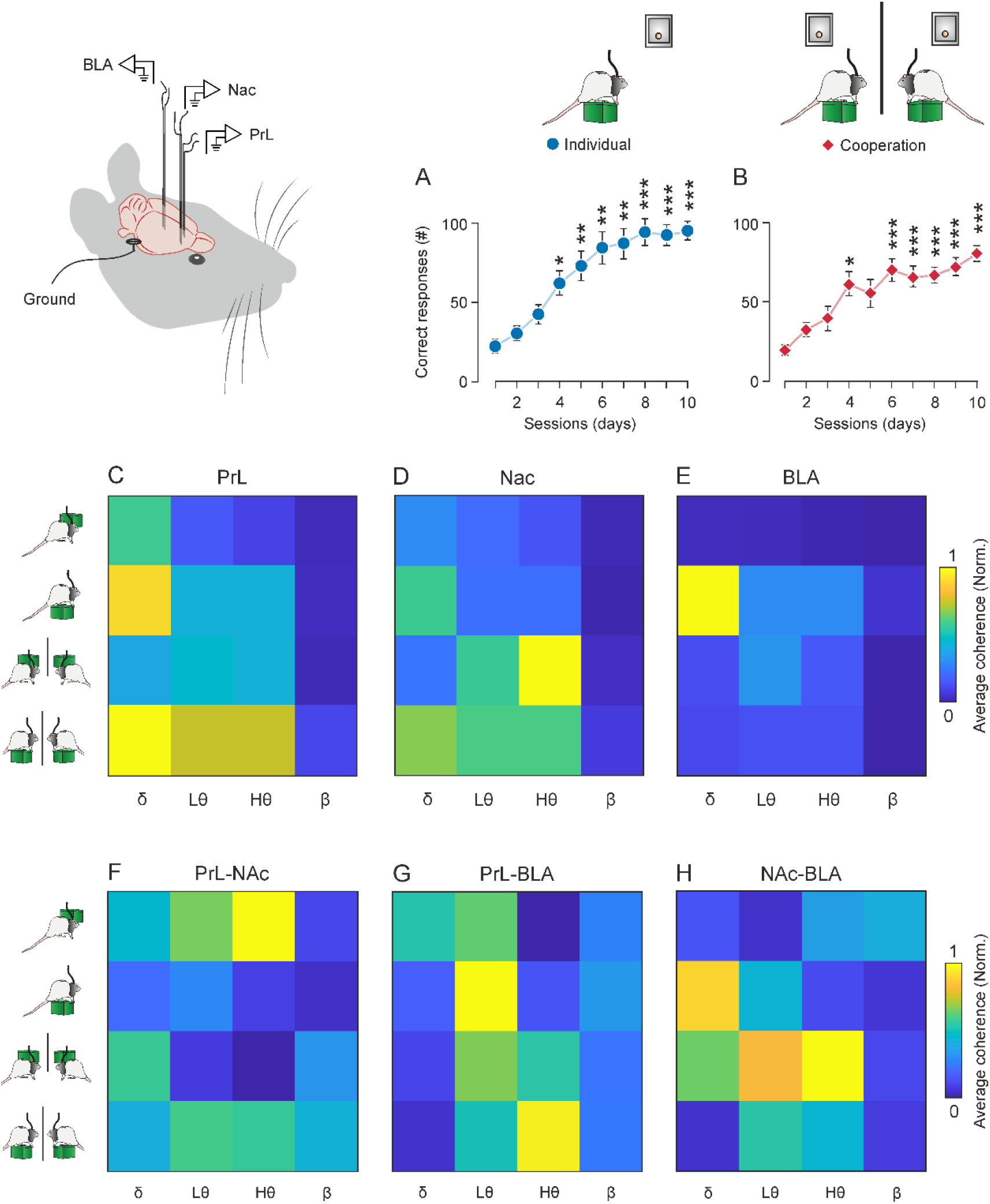
Cooperative task learning and general spectral power and coherence. A-B. Characteristics of the acquisition curves during the cooperation experiment. **A**, Phase I: Individual platform training. Rats (n = 10) were trained to individually climb onto a platform to obtain a pellet of food at a fixed 1:1 ratio. Rats were trained for up to 10 sessions. Compared with session 1, the number of correct responses increased significantly from session 4 on (RM-One-way ANOVA, S04, p< 0.05; S05, S06, S07, p < 0.01), showing an even greater increase in the last three sessions (RM-One-way ANOVA, S08, S09, S10, p < 0.001). **B**, Phase II: Cooperative training. Rats had to climb onto their respective platform and stay on it simultaneously for ≥ 0.5 s to mutually get a reward. Rats were trained for up to 10 sessions. Compared with session 1, the number of cooperation trials also increased significantly in session 4 (RM-One-way ANOVA, S04, p < 0.01), and from sessions 7 to 10 (RM-One-way ANOVA, S07, S08, S09, S10, p < 0.001). **C-E**, Visual summary of results found in LFP analysis from the 3 recording sites during both conditions and phases. **C**, Mean spectral power per band of LFPs in PrL cortex BEFORE- and ON-platform during individual and cooperative phases. Note the highest spectral power was found ON-platform during cooperation and ON-platform individually, although at a lower level. The theta band also presented high values during cooperation ON-platform. **D**, Same as in A for the NAc. Note that the highest spectral powers were found in high theta band BEFORE-platform and in delta ON-platform for the cooperative phase. **E**, Same as in A and B for the BLA. Note that the highest spectral power values were found in delta and theta bands when rats were individually ON-platform. **F-H**, Visual summary of results found in coherence analysis between the three recording sites during both conditions and phases. **F**, Mean coherence per band between LFPs recorded in PrL cortex and NAc, BEFORE- and ON-platform during individual and cooperative phases. Note the highest coherence values were found in the theta band BEFORE-platform during the individual phase and theta bands ON-platform during cooperation. Delta also showed high values before rats climbed onto the platform to cooperate. **G**, Same as in A for the PrL-BLA. Note that the highest coherence values were found in the low theta band when rats were individually ON-platform and in high theta when rats were cooperating ON-platform. **H**, Same as in A and B for NAc-BLA. Note that the highest coherence values were found in delta and theta bands BEFORE-platform. Delta also showed high coherence when rats were individually ON-platform.

During phase II, rats were trained to climb onto their respective platforms and stay on them simultaneously for at least 500 ms to mutually get a reward (a food pellet for each rat). As shown in the acquisition curve in Fig. 5B, rats learned to climb simultaneously onto the platform for the mutual reward, and the number of cooperation trials also increased significantly in session 4 (RM-One-way ANOVA, multiple comparisons, S04, p < 0.01), and from sessions 7 to 10 (RM-One-way ANOVA, multiple comparisons, S07, S08, S09, S10, p < 0.001).

To examine the oscillatory activity occurring in these brain areas during the two phases, 2-second epochs were selected from the continuous recordings obtained during two different situations: “BEFORE-platform”, comprising the 2 s before the animals climbed onto the platform, and “ON-platform”, which refers to the 2 s after the animal climbed on the platform.

Fig. 5C-E shows a visualization of normalized and averaged spectral-power results for all frequency bands during the individual and cooperative phase. In general, when rats were cooperating on the platform the PrL cortex presented higher power in delta, low theta, and high theta bands. These bands also showed increases in power when rats were individually on the platform, but to a lower degree. The highest powers for the NAc were also observed during the cooperative phase, when high theta increased before the climb onto the platform to cooperate, while delta and low theta did so after the climb onto the platform. The BLA showed less activation than the aforementioned areas, having the maximum activity when rats were individually on the platform, especially in the delta band.

To know more about the functional connectivity between the brain areas of interest, we calculated the LFP-LFP coherence in the frequency domain between electrodes from the three areas of interest: PrL-NAc, PrL-BLA, and NAc-BLA at different moments (BEFORE- and ON-platform) and for both phases (individual and cooperative). The epochs analyzed were the same as in the previous spectral power analysis (2-second epochs, NT = 70 per condition).

Fig. 5F-H shows a visualization of averaged LFP-LFP coherence results for all frequency bands and phases. In general, the connectivity between the PrL and NAc showed the highest coherence value in the theta band before the rats climbed onto the platform individually, followed by an increase in delta band before they climbed onto the platform to cooperate. During cooperation the connectivity in the delta band decreased, increasing in theta (low and high). For the PrL-BLA, the highest coherence values were found in the low theta band when rats were individually ON-platform and in high theta when rats were cooperating ON-platform. The NAc-BLA connectivity increased in delta and theta bands before the climb to cooperate. Delta also showed high coherence when rats were individually ON-platform.

#### Spectral power of PrL cortex, NAc, and BLA was higher before rats climbed onto the platform individually

Comparisons of the spectral power (Fig. 6A-C) and spectral dynamic analysis (spectrograms, Fig. 6D-I) of LFPs recorded from the two conditions —BEFORE- and ON-platform— during the individual phase revealed significantly higher power (jackknife estimates of the variance, p < 0.05) in the three selected brain areas before the animals climbed individually onto the platform as compared with the 2 s on the platform. In the PrL cortex, a significant increase in power was observed in the bandwidth in the range 8-11 Hz (Fig 6A; jackknife estimates of the variance, p < 0.05). The dynamic analysis represented in Fig. 6D indicates that the differences were particularly of note around 1 s after the rats climbed onto the platform, while the significant differences found (Fig. 6G) indicate higher activity for the 15-20 Hz band during the first second ON-platform and for the 10-20 Hz band 2 s after the climbing individually onto the platform. These differences were not detected by the comparison of average spectral power in Fig. 6A. In the NAc, the spectral power BEFORE-platform was significantly higher from 2 Hz to 6 Hz and from 9 Hz to 20 Hz (Fig. 6B), and the dynamic analysis confirmed these findings, revealing that the power values in the NAc were higher from 2 s to 1 s BEFORE-platform. The power values in the BLA were also significantly higher BEFORE-platform for the 2-5 Hz and 11-17 Hz bands (Fig. 6C). The spectrograms in Fig. 6F, I, revealed that the power increase in the range 2-5 Hz observed in Fig. 1C was greater around 1 s BEFORE-platform, while the differences found from 11 Hz to 17 Hz were greater 2 s and 500 ms BEFORE-platform.

**Figure 6.**
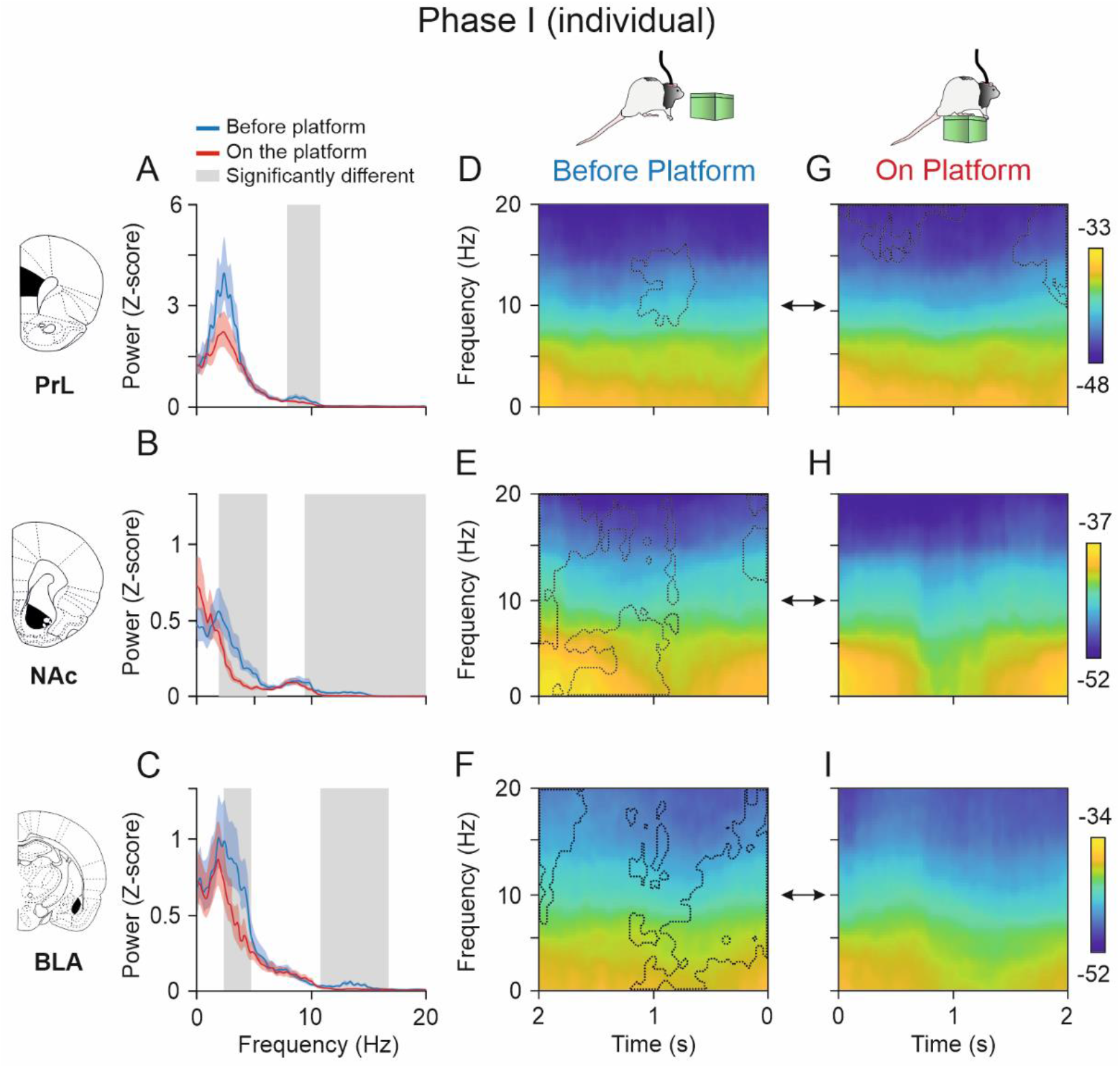
Spectral powers and multitaper spectrograms of LFPs recorded in the PrL cortex, NAc, and BLA areas during Phase I (individual). **A-C**, Spectral analysis for LFPs of 2-second epochs (NT = 70 per condition) acquired from five pairs of rats (n = 10) when they were individually ON-platform (red) compared with 2 s BEFORE-platform (blue). The lines represent the averaged spectrum of trials for each condition and the colored shaded areas the jackknife error bars. The gray shaded areas indicate the frequency ranges where the average spectral powers for each condition were significantly different. The spectral power in the delta band was significantly higher BEFORE-platform than ON-platform in the three areas for the lower frequencies. During the individual phase, the spectral power in the PrL cortex showed significantly higher power values in the theta band BEFORE-platform, while the NAc and BLA showed significantly higher spectral power in the high theta and beta bands. **D-I**, Multitaper spectrograms of the same epochs analyzed in **A-C** showing dynamic changes in LFP activities in the PrL cortex, NAc, and BLA areas 2 s BEFORE-(**D-F**) and 2 s ON-platform (**G-I**). Comparison of spectrograms for the two situations (BEFORE- and ON-). The dashed lines indicate the areas in which each spectrogram was significantly higher than the other (jackknife estimates of the variance, p < 0.05). In the PrL cortex (**D**, **G**), the spectral power of the theta band was significantly higher 1 s BEFORE-platform. ON-platform there were significantly higher power values for the beta band along second 1, and high theta and beta at the end of second 2. In the NAc (**E**, **H**), the spectral power BEFORE-platform was significantly higher than ON-platform, especially in the delta and low theta bands 2 s BEFORE-platform and high theta and beta 1.5 s BEFORE-the climbing onto the platform.

#### Spectral power of PrL cortex and NAc increased when rats climbed onto the platform to cooperate

Comparisons of the spectral power (Fig. 7A-C) and spectral dynamic analysis (spectrograms, Fig. 7D-I) of LFPs recorded for the two conditions —BEFORE- and ON-platform— during the cooperative phase showed a significant increase in the spectral power of PrL cortex in frequencies of 3-5 Hz, 7-10 Hz, and 14-19 Hz (jackknife estimates of the variance, p < 0.05) when rats were cooperating ON-platform compared with the seconds BEFORE-platform to cooperate (Fig. 7A). The dynamic analysis in Fig. 7D-G confirmed these findings and revealed that the spectral power was significantly higher, particularly at 0-10 Hz during the first second ON-platform and at 10-20 Hz after 500 ms ON-platform. (jackknife estimates of the variance, p < 0.05). The spectral power observed in the NAc (Fig. 7B) was significantly higher BEFORE-platform to cooperate at 7-11 Hz (jackknife estimates of the variance, p < 0.05). The comparison of multi-taper spectrograms in Fig. 7E-H showed that the predominant power BEFORE-platform was especially high for the band ranging between 7 Hz and 15 Hz 2 s and from 4 Hz to 15 Hz 1 s BEFORE-platform (jackknife estimates of the variance, p < 0.05). A significant increase in the power of the NAc at 3-4 Hz around 1.5 s after the climbing onto the platform was revealed in Fig. 7H. The spectral power of BLA was significantly higher at 3-20 Hz (Fig. 7C) and the spectrogram comparison shown in Fig. 7F-I indicated a significant decrease of power when rats were ON-platform for practically the whole time window analyzed.

**Figure 7.**
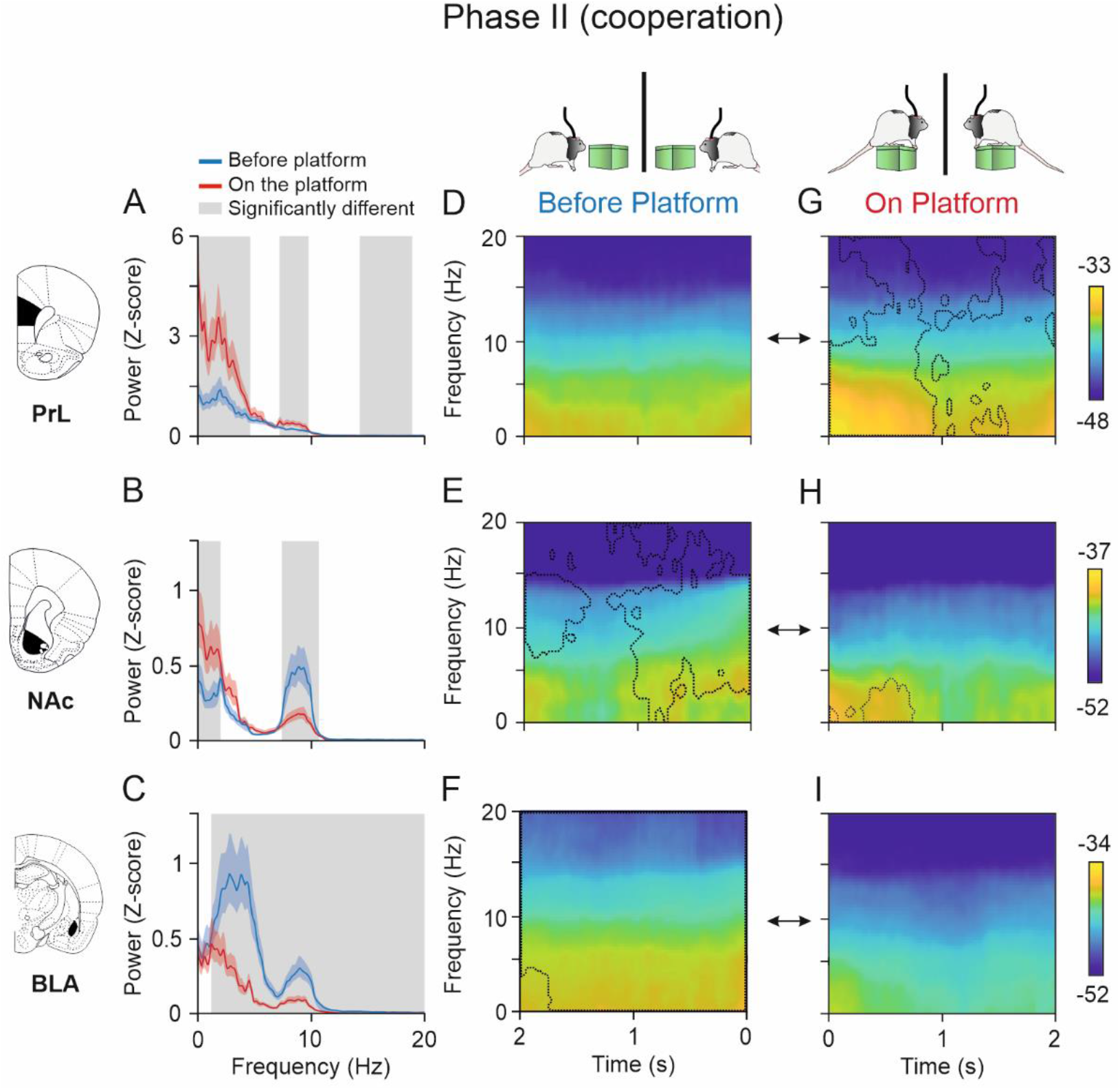
Spectral powers and multitaper spectrograms of LFPs recorded in the PrL cortex, NAc, and BLA areas during Phase II (cooperative). **A-C**, Same configuration as Fig. 6 but for the cooperative phase. The spectral power in the delta band was significantly higher BEFORE-platform than ON-platform in the three areas for the lower frequencies. Note that spectral power of the PrL cortex (**A**) was significantly higher when the rats were ON-platform in the delta, theta, and beta bands, while the spectral power observed in the NAc (**B**) was significantly higher in the delta band when rats were ON-platform, and higher in the theta band BEFORE-platform. The activity of the BLA (**C**) decreased significantly as rats climbed onto the platform, being significantly higher in all the frequency bands studied before the rats climbed onto the platform. **D-I**, Multitaper spectrograms of the same epochs analyzed in **A-C** showing dynamic changes in LFP activities in the PrL cortex, NAc, and BLA areas in the moments BEFORE-(**D-F**) and ON-platform (**G-I**). The spectrograms for the two situations, BEFORE- and ON-, were compared, and the dashed lines indicate the areas in which each spectrogram was significantly higher than the other (jackknife estimates of the variance, p < 0.05). The spectrograms indicated highest activity of delta and theta bands occurred within the first second ON-platform in the PrL cortex (**D**, **G**). In the NAc (**E**, **H**), the spectral power BEFORE-platform was significantly higher than ON-platform across almost the whole time window, while the activity in the delta band was higher within the first second ON-platform. In the BLA (**E**, **H**), the spectrograms showed higher power for almost the whole time window.

#### Coherence between PrL and NAc in the theta band increased before rats climbed individually onto the platform

After comparing the spectral powers from the structures under study, we compared the averaged spectral coherence during the two conditions (BEFORE- and ON-platform). After that, to obtain deeper knowledge about the time in which these changes in coherence were taking place for each area, we computed multitaper coherograms for each condition and compared them with the jackknife estimates of the variance method.

As shown in Fig. 8A, during the individual phase the coherence between the PrL cortex and the NAc was quite similar for the two different moments analyzed (BEFORE- and ON-platform) except for the range 8-11 Hz, where the coherence was significantly higher before rats climbed individually onto the platform (jackknife estimates of the variance, p < 0.05). The dynamic analysis illustrated in Fig. 8D-G revealed that the highest coherence magnitudes between the PrL cortex and the NAc were found BEFORE-platform, especially from 3 Hz to 4 Hz 1.5 s BEFORE- and from 13 Hz to 16 Hz, at around 1.5 s BEFORE-platform. The coherence between PrL cortex and the BLA was also similar for the two conditions (Fig. 8B), except around 6 Hz, which was significantly higher ON-platform (jackknife estimates of the variance, p < 0.05). The opposite happened at 10 Hz, where the coherence between the PrL cortex and the BLA increased BEFORE-platform. The coherograms in Fig. 8E-H revealed significantly higher coherence clusters BEFORE-platform, particularly 2 s BEFORE-in the range 6-9 Hz, and at 5 Hz and 15 Hz 1 s BEFORE-platform and 6 Hz to 9 Hz and 16 Hz 500 ms BEFORE-platform. The coherence between the NAc and the BLA was significantly higher when rats were individually ON-platform from 4 Hz to 6 Hz and 7 Hz to 10 Hz (Fig. 8C, jackknife estimates of the variance, p < 0.05) and the coherograms in Fig. 8F-I confirmed these differences, indicating significantly higher coherence from 3 Hz to 6 Hz 1 s after rats climbed individually onto the platform and from 7 Hz to 9 Hz 500 ms after the climbing ON-platform (jackknife estimates of the variance, p < 0.05). Some small clusters were significantly higher 500 ms before the climbing onto the platform at 8 Hz and 15 Hz.

**Figure 8.**
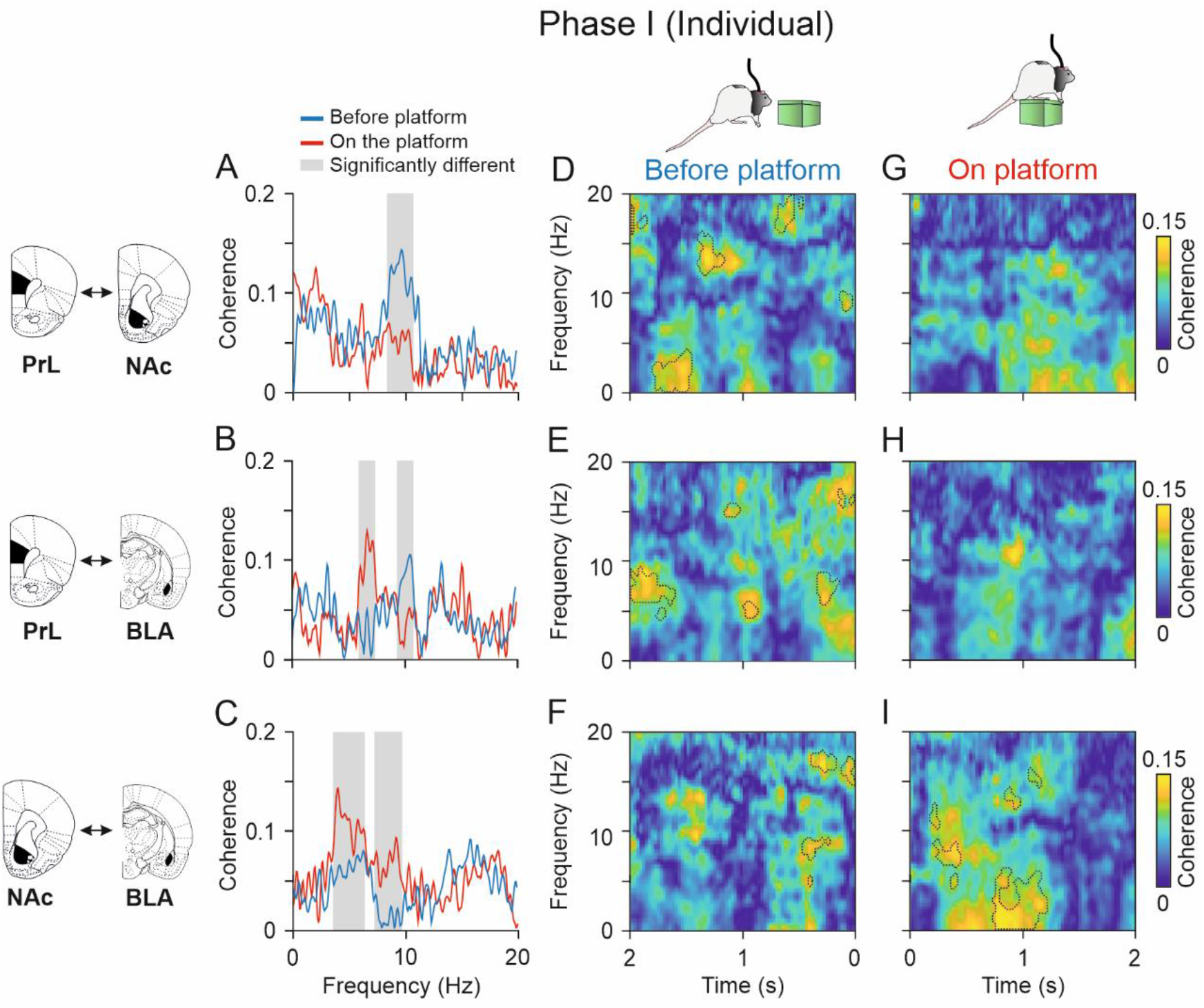
Functional connectivity between LFPs from electrodes located in different brain structures during Phase I (individual). **A-C**, Spectral coherence between the PrL cortex and NAc (**A**), PrL-BLA (**B**), and NAc-BLA (**C**) when rats were ON-platform (red) compared with 2 s BEFORE-platform (blue). The gray shaded areas indicate the frequency ranges where the average spectral powers for each condition were significantly different. **D-I,** time-frequency coherograms of the structures analyzed in **A-C** showing dynamic changes in phase coherence between the PrL cortex and NAc, PrL-BLA (jackknife estimates of the variance, p < 0.05), and NAc-BLA in the moments BEFORE-(**D-F**) and ON-platform (**G-I**). The coherograms for the two situations, BEFORE- and ON-, were compared, and the dashed lines indicate the areas in which each coherogram was significantly higher than the other (jackknife estimates of the variance, p < 0.05). Note that the coherence magnitude between PrL and NAc was higher BEFORE-platform than ON-platform in the theta and delta bands (jackknife estimates of the variance, p < 0.05). The coherence between PRL and BLA was higher in the low theta band when rats were ON-platform (jackknife estimates of the variance, p < 0.05), particularly 1 s after climbing, and in the high theta band 2 s and 500 ms BEFORE-platform. The coherence between NAc and BLA was significantly higher ON-platform (jackknife estimates of the variance, p < 0.05) and significantly higher in the delta and theta bands, particularly around 1 s after the climbing onto the platform.

#### The connectivity between the PrL cortex and the NAc in the high theta band increased when rats were cooperating on the platform

As shown in Fig. 9A, during the cooperative phase the coherence between the PrL cortex and the NAc was significantly higher when rats were cooperating ON-platform at 8-10 Hz and at 18-19 Hz (Fig. 9A, jackknife estimates of the variance, p < 0.05). This result was confirmed by the dynamic analysis illustrated in Fig. 9D-G, revealing the highest coherence magnitudes between the PrL cortex and the NAc 500 ms to 1 s after rats climbed ON-platform at 10-15 Hz, and another significantly higher cluster at 17-20 Hz right after they climbed ON-platform (jackknife estimates of the variance, p < 0.05). It is worth mentioning that the peak of coherence around 8-10 Hz observed during the individual phase before the climb onto the platform (Fig. 8A) disappeared during cooperation (Fig. 9A). In addition, coherence in this band during cooperation increased.

**Figure 9.**
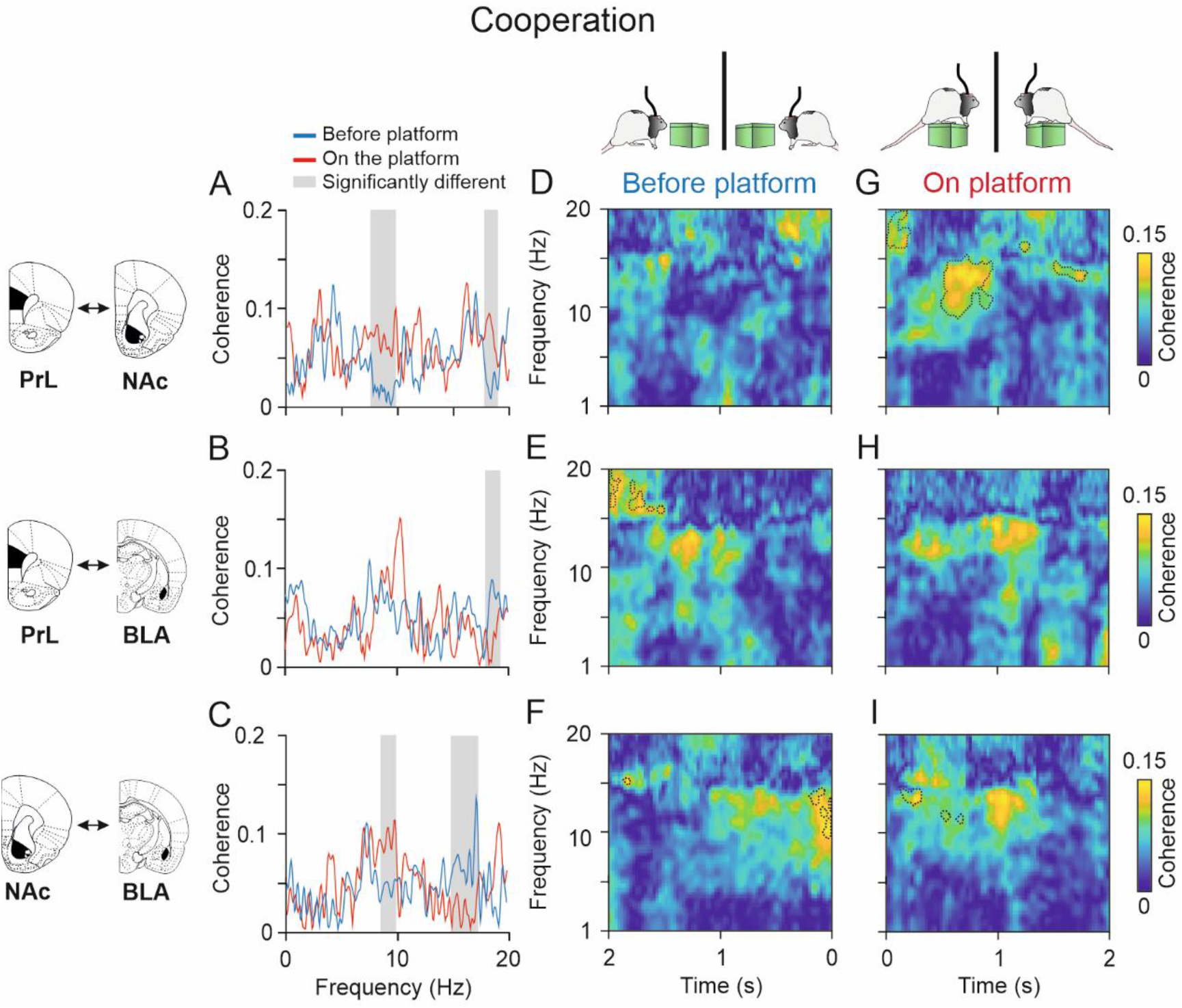
Functional connectivity between LFPs from electrodes located in different brain structures during Phase II (cooperative). **A-C**, Spectral coherence between the PrL cortex and NAc (**A**), PrL-BLA (B), and NAc-BLA (**C**) when rats were ON-platform (red) compared with 2 s BEFORE-platform (blue). The gray shaded areas indicate the frequency ranges where the average spectral powers for each condition were significantly different. **D-I,** time-frequency coherograms of the structures analyzed in **A-C** showing dynamic changes in phase coherence between the PrL cortex and NAc, PrL-BLA, and NAc-BLA in the moments BEFORE-(**D-F**) and ON-platform (**G-I**). The coherograms for the two situations, BEFORE- and ON-, were compared, and the dashed lines indicate the areas in which each coherogram was significantly higher than the other (jackknife estimates of the variance, p < 0.05). Note that the coherence magnitude between PrL and NAc was significantly higher ON-platform than BEFORE-platform in the theta and beta bands (jackknife estimates of the variance, p < 0.05). Note the significantly higher coherence cluster 0.5-1 s after rats climbed ON-platform at 10-15 Hz in the coherogram (**G**). The coherence between the NAc and the BLA was significantly higher when rats were cooperating ON-platform at 8-10 Hz and significantly higher BEFOREplatform at 15-17 Hz (jackknife estimates of the variance, p < 0.05).

The coherence between PrL cortex and the BLA was similar for the two conditions (Fig. 9B), except at 18-19 Hz. The peak coherence was found around 10 Hz, for the ON-platform condition, although the differences were not significant. The coherograms in Fig. 9E-H were very similar, revealing only small clusters of higher coherence at 17-20 Hz 2 s before the climbing onto the platform. The coherence between the NAc and the BLA was significantly higher when rats were cooperating ON-platform at 8-10 Hz (Fig. 9C) and significantly higher BEFORE-platform at 15-17 Hz (jackknife estimates of the variance, p < 0.05). The coherograms in Fig. 9F-I situated these differences in small clusters, showing the highest coherence at 9-14 Hz 200 ms BEFORE-platform and around 14 Hz 200 ms after the climbing onto the platform. The coherence between the PrL cortex and BLA and between NAc and BLA showed high coherence in the band at 10-15 Hz 1 s after the climbing ON-platform, but the differences with the BEFORE-platform conditions were not significant.

## Discussion

The present study convincingly shows that couples of rats are able to cooperate in an instrumental conditioning task for a mutual reinforcement. Beyond results reported in former studies (Lopuch and Popick, 2011; Carvalho et al., 2018; Conde-Moro et al., 2019), the present work identifies the specific PrL cell types engaged in this behavior, as well as the functional connectivity between the PrL with related brain regions (BLA, NAc) during cooperation.

### *Leader* and *follower* rats show different learning strategies and a distinct pattern of PrL activation

In this work, before the cooperation experiment, most *leader* rats presented low levels of anxiety, while most *follower* rats presented intermediate levels of anxiety (Fig. 3D-F). According to Herrero et al. (2006) and Venero et al. (2004), higher levels of basal anxiety in rodents are related to impaired learning and memory. This could explain why *follower* animals in our experiments —presenting higher trait-anxiety than *leaders*— had more difficulty in grasping the requirements to complete the cooperation trials. In this sense, *leader*s would likely be less anxious animals and present a better cognitive performance. The locomotor activity was similar for both *leaders* and *followers*, indicating that the differences observed in brain activity were not related to differences in locomotion. Regarding social competition, we hypothesized that *leader* rats would present higher levels of dominance than *followers*, but to our surprise, *follower*s scored higher in the social dominance index. These results were consistent with the Larrieu et al. (2017) study, in which animals classified as dominants also presented higher anxiety.

Results from the co-localization analysis in the PrL cortex indicated that *leader* rats participating in the cooperation phase presented higher c-FOS activation in glutamatergic neurons containing D1 receptors than *follower* ones. This activation was also higher than that presented by *leaders* and *followers* during the individual phase and by the control group, suggesting the involvement of these medium-spiny neurons in the acquisition of the cooperative task. Previous neuroanatomical studies found that most prefrontal cortex (PFC) glutamatergic neurons express either D1 or D2 receptors, rather than expressing both in the same neurons (Vincent et al., 1993; Gaspar et al., 1995; Santana et al., 2009). This anatomical differentiation separates PFC dopaminergic neurons in two populations, which would project to different subcortical areas.

We also fund higher c-FOS activation of glutamatergic cells containing D2 receptors in the cooperation group than in the control and the individual groups (that did not participate in the cooperation phase), although the differences between *leaders* and *followers* were not significant, suggesting that these cells could also play a role in the cooperation process, requiring further analysis.

### PrL and related subcortical circuits (NAc and BLA) are involved in the cooperative acquisition of an instrumental conditioning task

After showing the involvement of the PrL cortex in cooperation and identifying cell types that increased their activation during task performance, we recorded LFPs from other subcortical structures, such as the NAc and the BLA, known to receive and send projections to PrL neurons and to have an important role in social behaviors.

Rats completed the cooperation task successfully, but rats’ strategies to cooperate were more homogeneous than in previous experiments (Conde-Moro et al., 2019), and there was not a clear difference between *leaders* and *followers*. This might be due to the adaptations of the cooperation experiment or to differences between rat batches, as some might contain less anxious rats, which, as mentioned before, should present fewer cognitive impairments and thus show better performance in the task (Venero et al., 2004; Herrero et al., 2006).

The analysis of the averaged spectral power and time-frequency decomposition showed that the highest spectral power was found in the PrL (delta band) when rats were cooperating on the platform. This increase was greater during the first second after the climbing onto the platform. When rats were cooperating on the platform, the power in low and high theta bands also increased significantly compared with moments of resting individually on the platform or before climbing onto it. These findings support those found in our previous work (Conde-Moro et al., 2019) suggesting the involvement of the PrL cortex in the acquisition of a cooperative task. LFPs recorded in the NAc showed interesting power dynamics as well. Delta and low theta ranges were selectively more active during the first second of cooperation, while the high theta band had a considerable dip at this same moment. As previously noted, dopaminergic neurons located in the rat PrL cortex project to the NAc (Ongür and Price, 2000) and to other subcortical structures that, in conjunction, can modulate social behavior (Grossman, 2013; Gunaydin et al., 2014). According to Goto and Grace (2015), a dopamine release from the PFC can modulate the activity of the NAc in goal-directed behaviors. The co-localization results in this study showed that the activity of D1-containing cells in the PrL is increased in *leader* rats during cooperation. Jenni et al. (2017) found that dopamine in PFC D1 receptors reinforces responses, providing larger rewards through connections to the NAc, while dopamine on PFC D2 receptors facilitates adjustments in decision-making through connections to the BLA. Following this argument, it would be safe to assume that the D1 neurons that were more active during the cooperation task in our rats are those likely projecting to the NAc, rather than to the BLA.

Additionally, in the connectivity analysis, we observed that when rats were cooperating on the platform, there was higher phase coherence between the LFPs recorded in the PrL cortex and the NAc in the 9-15 Hz frequency range (high theta) than before they climbed onto the platform, especially 1.5-1 s on the platform. BEFORE-platform, the highest coherence was found in the delta and theta bands 0.5-1 s before the climbing onto the platform. These findings add more evidence to suggest an involvement of populations of cells in the NAc in the decision of climbing onto the platform to cooperate or in the prediction of a mutual reward by cooperating with a conspecific.

According to Liu et al. (1994), Haber and McFarland (1999), and Goto and Grace (2015), the NAc could act as a gatekeeper, controlling what information is important enough to get access to basal ganglia nuclei, such as the amygdala. In this work, the highest power found in the BLA was in the delta, and low and high theta bands, when rats were individually on the platform. The connectivity analysis also indicated higher levels of coherence between the NAc and the BLA when rats were individually on the platform. The highest coherence during the cooperation phase was found between the NAc and the BLA in the theta band before the climbing onto the platform, which is consistent with the accentuated decrease in BLA power in all the bands and across the whole time window observed during the cooperation phase. It is known that the BLA sends strong projections to the PRL cortex and the NAc (Groenewegen et al., 1990; Haber et al., 1995), structures known to modulate emotional and motivational processing, including the motivational aspects of predicting cues (Davis, 1922; Everitt et al., 2000). Thus, we consider that the BLA might be involved in the prediction and anticipation of the task outcome.

In conclusion, the present study identified specific neuronal types from the PrL cortex engaged in the acquisition of a cooperative task. Rats leading the cooperation trials presented an increased c-FOS activation of D1-containing neurons in the PrL during cooperation. The PrL and NAc electrical activity (LFPs) increased when rats were cooperating, while the BLA activity increased before cooperation. The PrL and NAc showed increased functional connectivity at the moment of cooperation on the platform, whereas during the individual phase the highest connectivity was found before the animals climbed individually onto the platform. Additionally, we consider that the changes in specific cells and electrical activity observed in this work were due to the cooperative aspect of the task and not to the rewarding effect that other task requirements could have for the rats (such as climbing onto the platform) or for the mere social component of it (i.e., being in the apparatus with a cagemate), as all of these conditions were also met during the individual phase.

## Acknowledgements

This work was supported by grants PY18-823 and BIO-122 from the Spanish Junta de Andalucía, and the Swiss National Science Foundation (NCCR Synapsy 51NF40-158776 and −185897). We thank Pier L. Giussani, José A. Santos-Naharro and Olivia Zanoletti for their excellent technical assistance. We also thank Mr. Roger Churchill for his careful revision of the final version of the manuscript.

## Extended data

**Figure 1-1.**
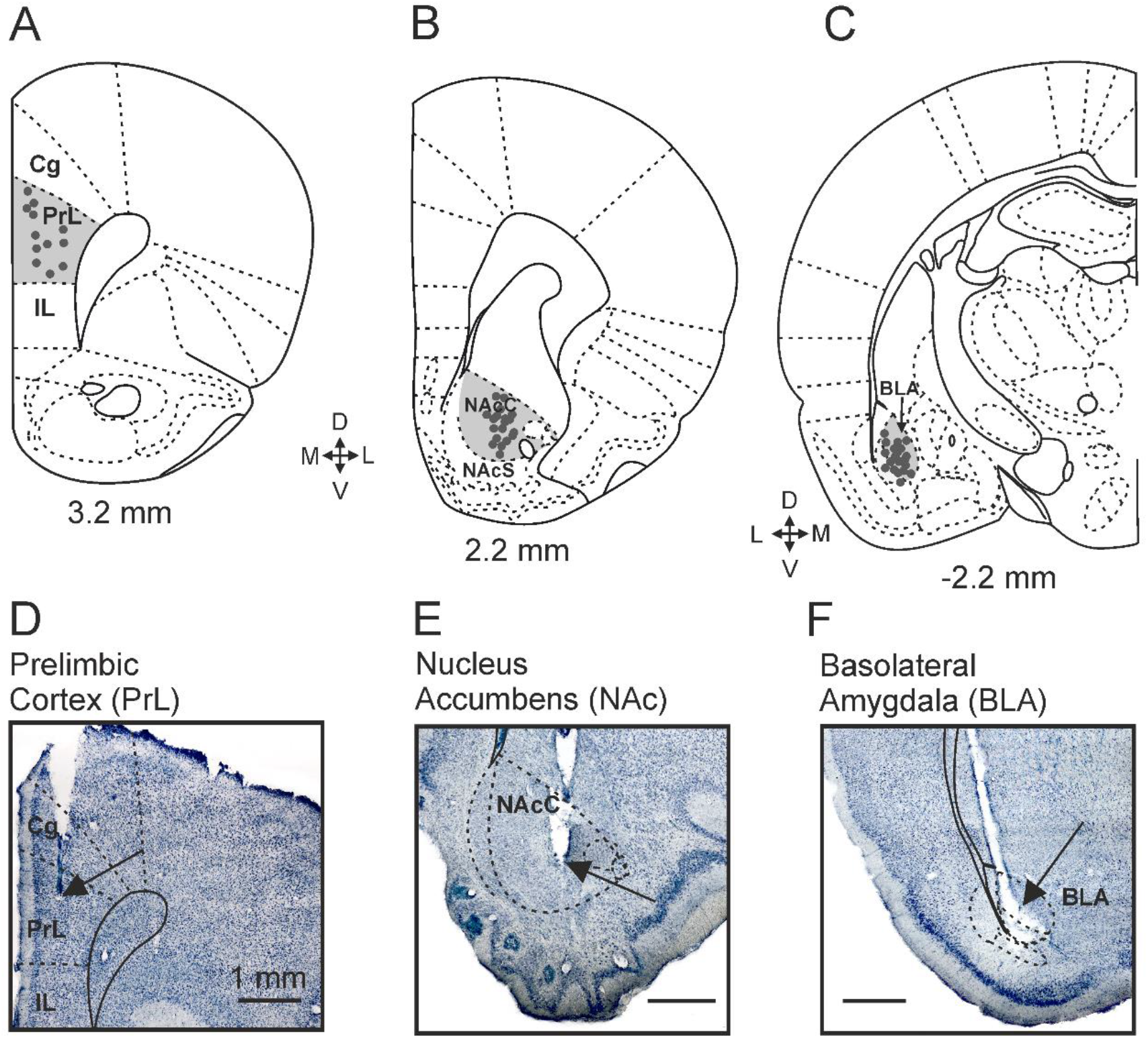
**A-C.** Diagrams representing the final locations of each electrode in the three areas recorded. **A**, each rat was implanted with two sets of recording electrodes aimed at the right PrL cortex (3.24 anterior and 0.5 mm lateral to bregma, and 2.5 mm from the surface). **B**, each rat was implanted with two sets of recording electrodes aimed at the right NAc (2.2 mm anterior and 1.5 mm lateral to bregma, and 6.5 mm from the surface). **C**, a third set of recording electrodes was aimed at the left BLA (2.28 posterior and 5 mm lateral to bregma, and 7.5 mm from the surface). **D-I**, photomicrographs of the brain regions of interest showing the final location of the recording electrodes. The tissue was dyed following the Nissl technique. The arrows point to the scar left by the electrode in the tissue.

**Figure 2-1.**
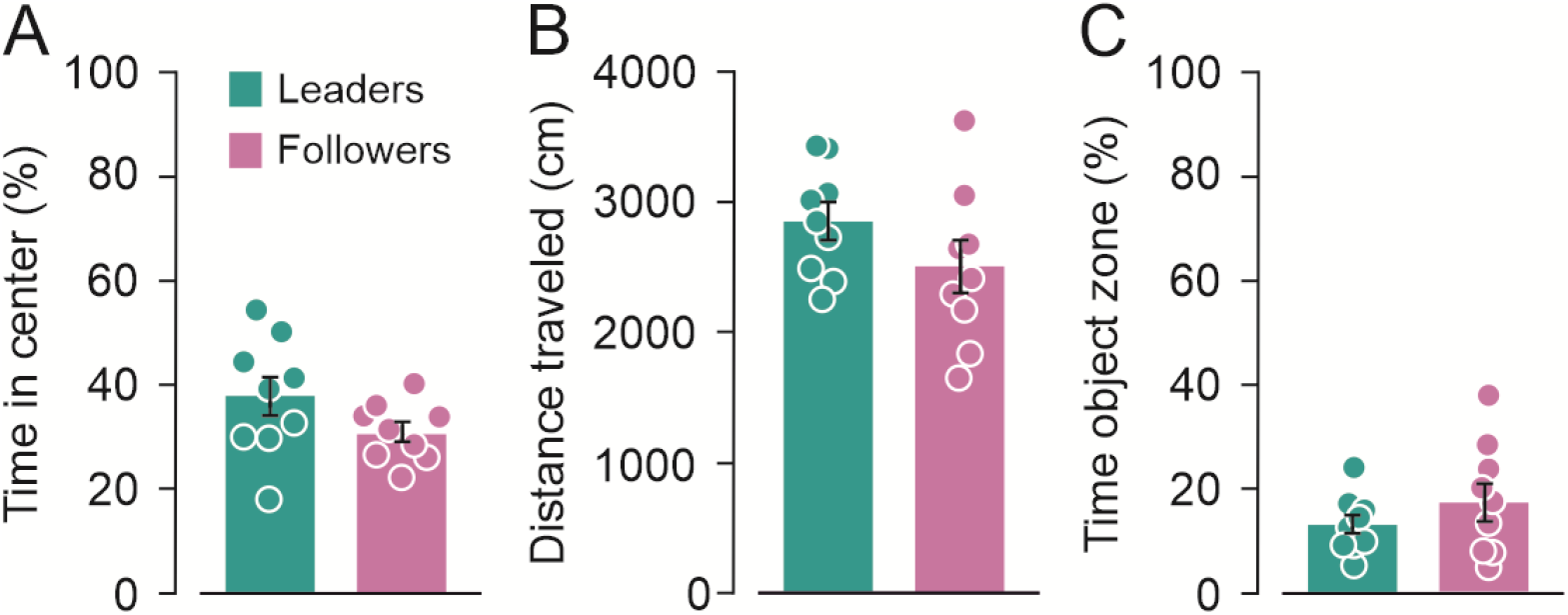
Open field test (groups 2 and 3). **A**, percentage of time spent in the center and border areas of the OFT. Although *leader* rats spent more time in the open arms —more anxiogenic— than *followers*, there was high variability among subjects, and the difference between groups was not significant (One-way ANOVA, F = 2.58, p = 0.12). **B**, distance traveled. Again, *leader* rats, in general, traveled more distance than *followers*, but the differences were not significant (One-way ANOVA, F = 2.58, p = 0.12). **C**, time spent in the novel object zone. Non-significant differences were found between the percentage of time that *leader* and *follower* rats spent in the object zone (One-way ANOVA, F = 1.10, p = 0.30).

**Figure 4-1.**
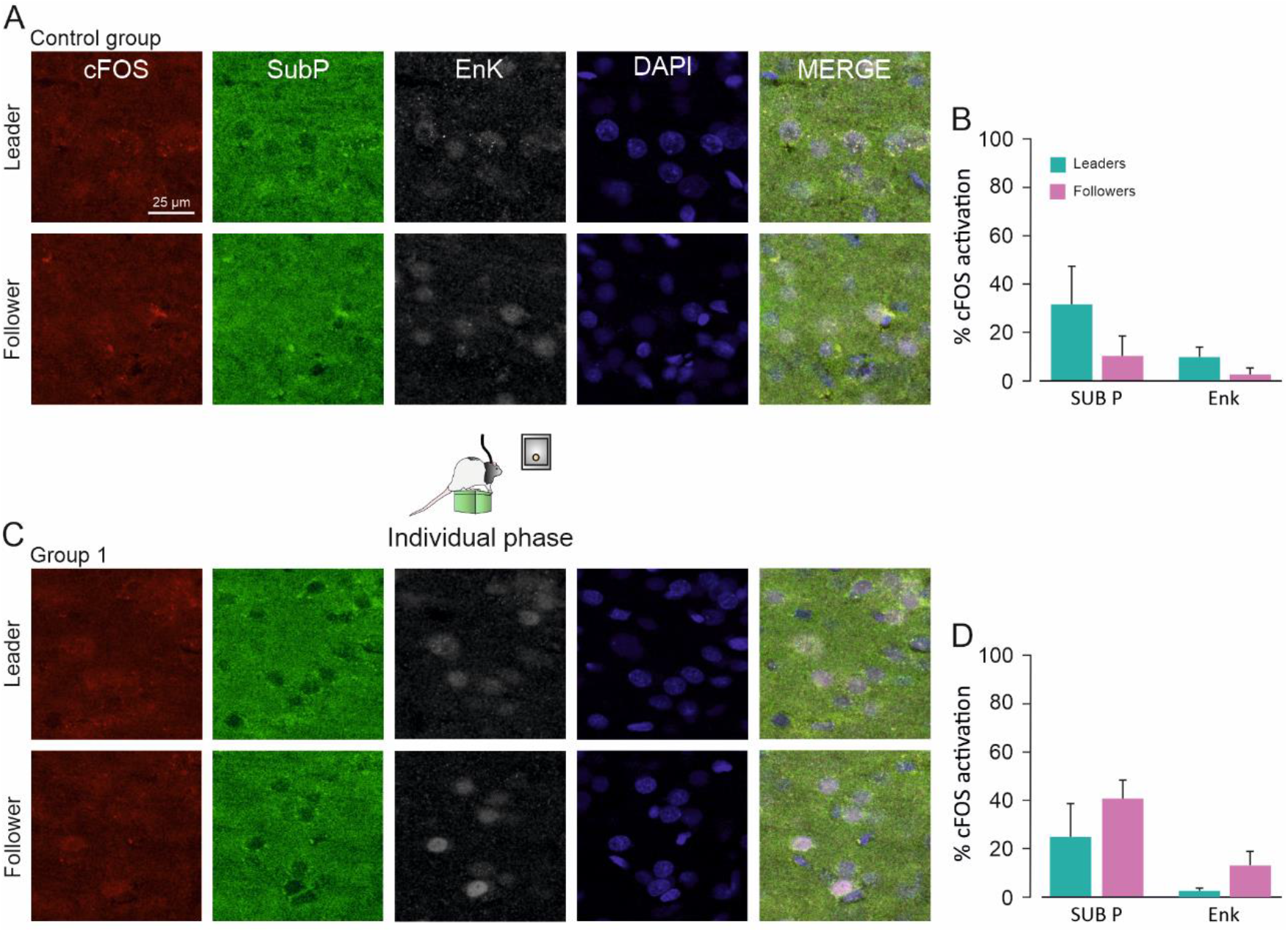
c-FOS expression in D1- and D2-containing cells in the PrL cortex of the individual (group 1) and control group. **A, C,** Same configuration as in Fig. 4.A-C. The photomicrographs corresponding to the control group, which did not participate in the experiments, are shown in A. The photomicrographs in **C** correspond to Group 1, in which rats were trained for the individual phase until reaching the criterion. **B**, **D**, The percentage of c-FOS activation in each group of cells was quantified. *Predicted follower* rats presented higher percentages of c-FOS expression than *predicted leaders*, but non-significant differences were found both in the level of activation of D2 cells during the individual phase (**D**, One-way ANOVA, F = 0.68, p = 0.42) and in the control group (**B,** One-way ANOVA on ranks, H = 0.02, p = 0.93). Non-significant differences were found both in the level of activation of D2 cells for *predicted leaders* and *predicted followers* during the individual phase (**D**, One-way ANOVA on ranks, H = 2.41, p = 0.11) and in the control group (**B**, One-way ANOVA, F = 1.85, p = 0.20).

**Figure 4-2.**
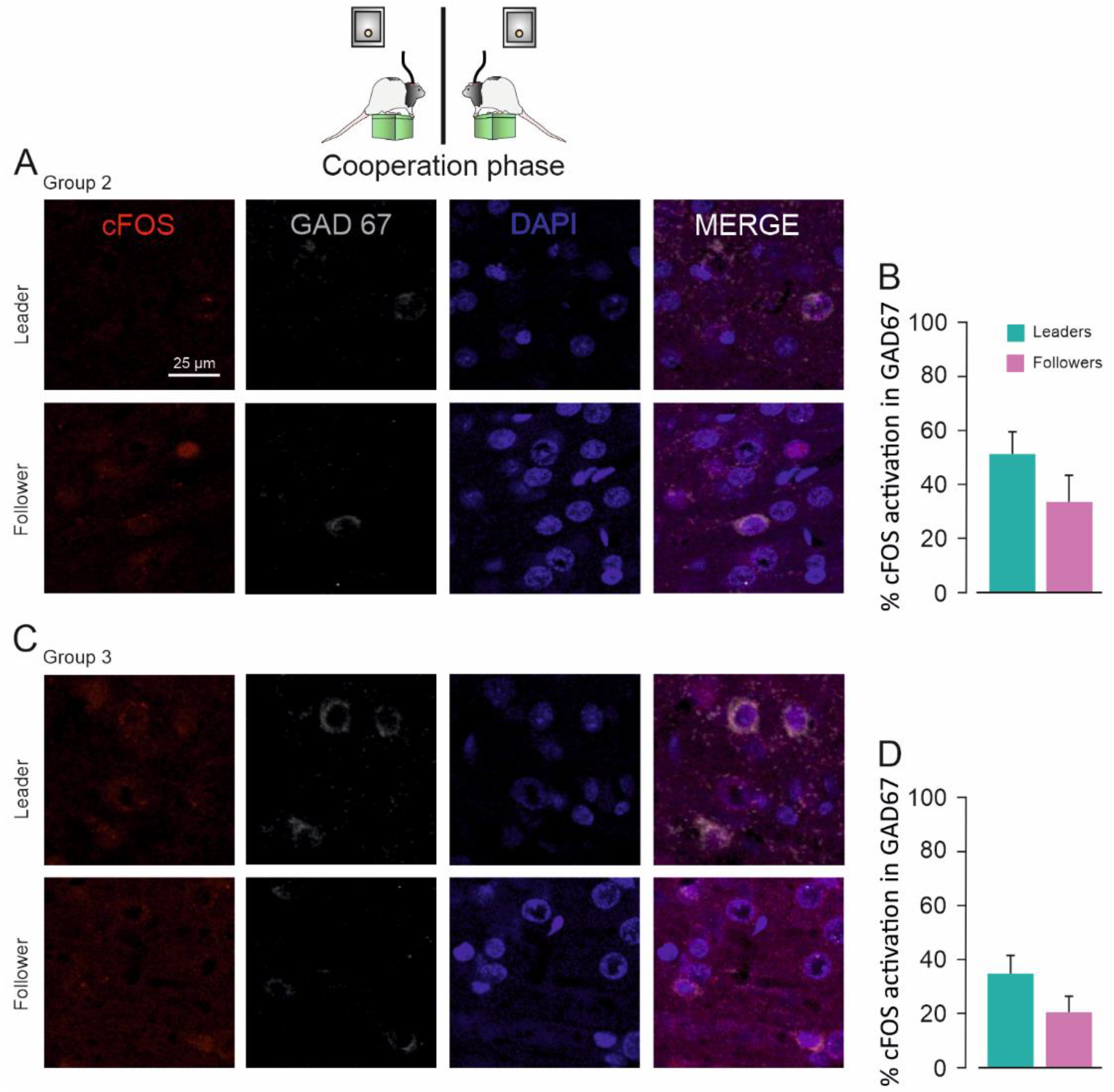
c-FOS expression in PrL GABAergic cells of cooperation groups (groups 2 and 3). Coronal sections from the four groups of animals were labeled for c-FOS (red channel), DAPI (blue channel), and mouse anti-GAD67 (gray channel) antibody for staining GABAergic cells of the PrL cortex. The photomicrographs corresponding to group 2, in which rats were trained to cooperate until reaching the criterion, are shown in **A**. The photomicrographs in **C** correspond to group 3, which completed the 10 days of the task. The percentage of c-FOS activation in GABAergic cells was quantified (**B-D**). Although *follower* rats in groups 2 and 3 showed a tendency for activation of GABAergic cells greater than that of *leaders*, no significant differences were found (**B**, One-way ANOVA, F =2.50, p = 0.12) and 3 (**D**, One-way ANOVA, F = 1.62, p = 0.22).

**Figure 4-3.**
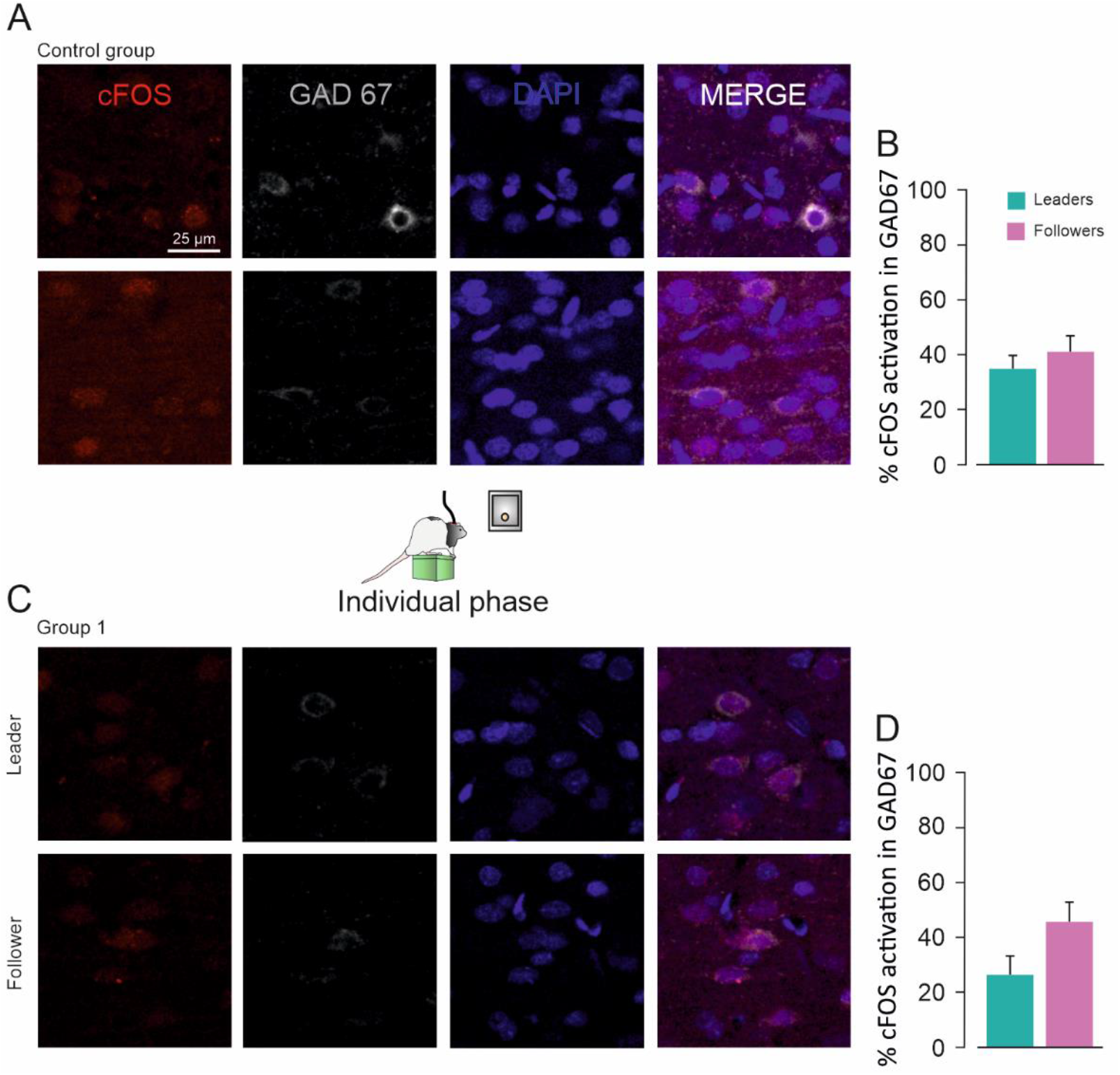
c-FOS expression in PrL GABAergic cells of control group and group 1. Coronal brain sections were labeled for c-FOS (red channel), DAPI (blue channel), and mouse anti-GAD67 (gray channel) antibody for staining GABAergic cells of the PrL cortex. The photomicrographs corresponding to the control group, which did not participate in the experiments, are shown in **A**. The photomicrographs in **B** correspond to group 1, which performed the individual task until reaching criterion. The percentage of c-FOS activation in GABAergic cells was quantified (**B-D**). Although *predicted follower* rats in group 1 showed a tendency for higher activation of GABAergic cells, non-significant differences were found between *predicted leaders* and *predicted followers* in both of these groups (**B**, One-way ANOVA, F = 2.15, p = 0.16; **D**, One-way ANOVA, F = 0.49, p = 0 .49).

## Notes

**Conflict of interest**: The authors declare no competing financial interest.

### Competing Interest Statement

The authors have declared no competing interest.

## References

1. Adolphs R (2010) What does the amygdala contribute to social cognition? Ann N Y Acad Sci 1191:42–61. doi: 10.1111/j.1749-6632.2010.05445.x

2. Bartal IB-A, Decety J, Mason P (2011) Empathy and Pro-Social Behavior in Rats. Science 334:1427–1430. doi: 10.1126/science.1210789

3. Bartal IB-A, Breton JM, Sheng H, Long KL, Chen S, Halliday A, Kenney JW, Wheeler AL, Frankland P, Shilyansky C, Deisseroth K, Keltner D, Kaufer D (2021) Neural correlates of ingroup bias for prosociality in rats. Elife 10:e65582. doi: 10.7554/eLife.65582

4. Bickart KC, Dickerson BC, Barrett LF (2014) The amygdala as a hub in brain networks that support social life. Neuropsychologia 63:235–248. doi: 10.1016/j.neuropsychologia.2014.08.013

5. Bokil H, Andrews P, Kulkarni JE, Mehta S, Mitra PP (2010) Chronux: A platform for analyzing neural signals. J Neurosci Meth 192:146–151. doi: 10.1016/j.jneumeth.2010.06.020

6. Bokil H, Purpura K, Schoffelen JM, Thomson D, Mitra P (2007) Comparing spectra and coherences for groups of unequal size. J Neurosci Meth 159:337–345. doi: 10.1016/j.jneumeth.2006.07.011

7. Carvalho LC, Dos Santos L, Regaço A, Barbosa TB, da Silva RF, de Souza D das G, Sandaker I (2018) Cooperative responding in rats maintained by fixed- and variable-ratio schedules. J Exp Anal Behav 110:105–126. doi: 10.1002/jeab.457

8. Chalmeau R, Visalberghi E, Gallo A (1997) Capuchin monkeys, Cebus apella, fail to understand a cooperative task. Anim Behav 54:1215–1225. doi: 10.1006/anbe.1997.0517

9. Conde-Moro, AR, Rocha-Almeida, F, Sánchez-Campusano, R, Delgado-García, JM, Gruart, A (2019). The activity of the prelimbic cortex in rats is enhanced during the cooperative acquisition of an instrumental learning task. Prog Neurobiol, 183:101692. doi: 10.1016/j.pneurobio.2019.101692

10. Cordero MI, Sandi C (2007) Stress Amplifies Memory for Social Hierarchy. Front Neurosci 1:175–184. doi: 10.3389/neuro.01.1.1.013.2007

11. Costa DF, Moita MA, Márquez C (2021) Novel competition test for food rewards reveals stable dominance status in adult male rats. Sci Rep 11:14599. doi: 10.1038/s41598-021-93818-0

12. Crawford MP (1937) The cooperative solving of problems by young chimpanzees. Comparative Psychology Monographs, 14, 1–88. doi: 10.1080/00224545.1941.9714077

13. Crawford MP (1941) The Cooperative Solving by Chimpanzees of Problems Requiring Serial Responses to Color Cues. J Soc Psychol 13:259–280. doi: 10.1080/00224545.1941.9714077

14. Davis M (1992) The role of the amygdala in fear and anxiety. Annu Rev Neurosci, 15:353–375. doi: 10.1146/annurev.ne.15.030192.002033

15. Decety J, Svetlova M (2012) Putting together phylogenetic and ontogenetic perspectives on empathy. Dev Cogn Neurosci 2:1–24. doi: 10.1016/j.dcn.2011.05.003

16. Eisenberg N, Miller PA (1987) The relation of empathy to prosocial and related behaviors. Psychol Bull 101:91–119. Doi: 10.1037/0033-2909.101.1.91

17. Everitt BJ, Robbins TW (2000). Second-order schedules of drug reinforcement in rats and monkeys: measurement of reinforcing efficacy and drug-seeking behaviour. Psychopharmacology 153:17–30. doi: 10.1007/s002130000566

18. Felix-Ortiz AC, Tye KM (2014) Amygdala inputs to the ventral hippocampus bidirectionally modulate social behavior. J Neurosci 34:586–595. doi: 10.1523/JNEUROSCI.4257-13.2014

19. Gabbott PL, Warner T, Jays PRL, Salway P, Busby SJ (2005) Prefrontal cortex in the rat: Projections to subcortical autonomic, motor, and limbic centers. J Comp Neurol 492:145–177. doi: 10.1002/cne.20738

20. Gaspar P, Bloch B, Le Moine C (1995) D1 and D2 receptor gene expression in the rat frontal cortex: cellular localization in different classes of efferent neurons. Eur J Neurosci 7:1050–1063. doi: 10.1111/j.1460-9568.1995.tb01092.x

21. Goto Y, Grace AA (2005) Dopaminergic modulation of limbic and cortical drive of nucleus accumbens in goal-directed behavior. Nat Neurosci 8:805–812. doi: 10.1038/nn1471

22. Groenewegen HJ, Berendse HW (1990). Connections of the subthalamic nucleus with ventral striatopallidal parts of the basal ganglia in the rat. J Comp Neurol 294:607–622. doi: 10.1016/j.stemcr.2021.11.014

23. Grossmann T (2013) The role of medial prefrontal cortex in early social cognition. Front Hum Neurosci 7:1–6. doi: 10.3389/fnhum.2013.00340

24. Gunaydin LA, Grosenick L, Finkelstein JC, Kauvar IV, Fenno LE, Adhikari A, Lammel S, Mirzabekov JJ, Airan RD, Zalocusky KA, Tye KM, Anikeeva P, Malenka RC, Deisseroth K (2014) Natural neural projection dynamics underlying social behavior. Cell 157:1535–1551. doi: 10.1016/j.cell.2014.05.017

25. Haber SN, Kunishio K, Mizobuchi M, Lynd-Balta E (1995) The orbital and medial prefrontal circuit through the primate basal ganglia. J Neurosci 15:4851–4867. doi: 10.1523/JNEUROSCI.15-07-04851.1995

26. Haber SN, McFarland NR (1999) The concept of the ventral striatum in nonhuman primates. Ann NY Acad Sci 877(1), 33–48. doi: 10.1111/j.1749-6632.1999.tb09259.x

27. Hernandez-Lallement J, Van Wingerden M, Marx C, Srejic M, Kalenscher T (2015) Rats prefer mutual rewards in a prosocial choice task. Front Neurosci 9:1–9. doi: 10.3389/fnins.2014.00443

28. Herrero AI, Sandi C, Venero C (2006) Individual differences in anxiety trait are related to spatial learning abilities and hippocampal expression of mineralocorticoid receptors. Neurobiol Learn Mem 86:150–159. doi: 10.1016/j.nlm.2006.02.001

29. Hirata S, Fuwa K (2007) Chimpanzees (Pan troglodytes) learn to act with other individuals in a cooperative task. Primates 48:13–21. doi: 10.1007/s10329-006-0022-1

30. Hollis F, Van Der Kooij MA, Zanoletti O, Lozano L, Cantó C, Sandi C (2015) Mitochondrial function in the brain links anxiety with social subordination. Proc Natl Acad Sci U S A 112:15486–15491. doi: 10.1073/pnas.1512653112

31. Hoover WB, Vertes RP (2007) Anatomical analysis of afferent projections to the medial prefrontal cortex in the rat. Brain Struct Funct 212:149–179. doi: 10.1007/s00429-007-0150-4

32. Kingsbury L, Huang S, Wang J, Gu K, Golshani P, Wu YE, Hong W (2019) Correlated Neural Activity and Encoding of Behavior across Brains of Socially Interacting Animals. Cell 178:429–446.e16. doi: 10.1016/j.cell.2019.05.022

33. Larrieu T, Cherix A, Duque A, Rodrigues J, Lei H, Gruetter R, Sandi C (2017) Hierarchical Status Predicts Behavioral Vulnerability and Nucleus Accumbens Metabolic Profile Following Chronic Social Defeat Stress. Curr Biol 27:2202–2210.e4. doi: 10.1016/j.cub.2017.06.027

34. Lee E, Rhim I, Lee JW, Ghim J-W, Lee S, Kim E, Jung MW (2016) Enhanced Neuronal Activity in the Medial Prefrontal Cortex during Social Approach Behavior. J Neurosci 36:6926–6936. doi: 10.1523/JNEUROSCI.0307-16.2016

35. Liu L, Shen RY, Kapatos G, Chiodo LA (1994) Dopamine neuron membrane physiology: characterization of the transient outward current (IA) and demonstration of a common signal transduction pathway for IA and IK. Synapse, 17:230–240. doi: 10.1002/syn.890170404

36. Łopuch S, Popik P (2011) Cooperative behavior of laboratory rats (Rattus norvegicus) in an instrumental task. J Comp Psychol 125:250–253. doi: 10.1037/a0021532

37. Márquez C, Rennie SM, Costa DF, Moita MA (2015) Prosocial Choice in Rats Depends on Food-Seeking Behavior Displayed by Recipients. Curr Biol 25:1736–1745. doi: 10.1016/j.cub.2015.05.018

38. Mendres KA, de Waal FBM (2000) Capuchins do cooperate: the advantage of an intuitive task. Anim Behav 60:523–529. doi: 10.1006/anbe.2000.1512

39. Minami C, Shimizu T, Mitani A (2017) Neural activity in the prelimbic and infralimbic cortices of freely moving rats during social interaction: Effect of isolation rearing. PLoS One 12:e0176740. doi: 10.1371/journal.pone.0176740

40. Mitra PP, Bokil H (2008) Observed Brain Dynamics. New York, NY, USA: Oxford University Press.

41. Murugan M, Jang HJ, Park M, Miller EM, Cox J, Taliaferro JP, Parker NF, Bhave V, Hur H, Liang Y, Nectow AR, Pillow JW, Witten IB (2017) Combined Social and Spatial Coding in a Descending Projection from the Prefrontal Cortex. Cell 171:1663–1677.e16. doi: 10.1016/j.cell.2017.11.002

42. Nessler JA, Gilliland SJ (2009) Interpersonal synchronization during side by side treadmill walking is influenced by leg length differential and altered sensory feedback. Hum Mov Sci 28:772–785. doi: 10.1016/j.cell.2019.05.022

43. Ongür D, Price JL (2000) The organization of networks within the orbital and medial prefrontal cortex of rats, monkeys and humans. Cereb Cortex 10:206–219. doi: 10.1093/cercor/10.3.206

44. Paxinos, G, Watson C (2007) The Rat Brain in Stereotaxic Coordinates. San Diego, CA, USA: Academic Press.

45. Petit O, Desportes C, Thierry B (1992) Differential Probability of “Coproduction” in Two Species of Macaque (Macaca tonkeana, M. mulatta). Ethology 90:107–120. doi: 10.1111/j.1439-0310.1992.tb00825.x

46. Raihani NJ, Bshary R (2011) Resolving the iterated prisoner’s dilemma: theory and reality. J Evol Biol 24:1628–1639. doi: 10.1111/j.1420-9101.2011.02307.x

47. Rutte C, Taborsky M (2007) Generalized reciprocity in rats. PLoS Biol 5:1421–1425. doi: 10.1371/journal.pbio.0050196

48. Santana N, Mengod G, Artigas F (2009) Quantitative analysis of the expression of dopamine D1 and D2 receptors in pyramidal and GABAergic neurons of the rat prefrontal cortex. Cereb Cortex 19:849–860. doi: 10.1093/cercor/bhn134

49. Sato N, Tan L, Tate K, Okada M (2015) Rats demonstrate helping behavior toward a soaked conspecific. Anim Cogn. doi: 10.1007/s10071-015-0872-2

50. Schuster R, Perelberg A (2004) Why cooperate? An economic perspective is not enough. Behav Processes 66:261–277. doi: 10.1016/j.beproc.2004.03.008

51. Sesack SR, Deutch AY, Roth RH, Bunney BS (1989) Topographical organization of the efferent projections of the medial prefrontal cortex in the rat: An anterograde tract-tracing study with Phaseolus vulgaris leucoagglutinin. J Comp Neurol 290:213–242. doi: 10.1002/cne.902900205

52. Skinner BF (1962). Two “synthetic social relations”. J Exp Anal Behav 5(4), 531. doi: 10.1901/jeab.1962.5-531

53. Thomson DJ (1982) Spectrum estimation and harmonic analysis. Pr Inst Electr Elect, 70:1055–1096. doi: 10.1109/PROC.1982.12433

54. Venero C, Tilling T, Hermans-Borgmeyer I, Herrero AI, Schachner M, Sandi C (2004) Water maze learning and forebrain mRNA expression of the neural cell adhesion molecule L1. J Neurosci Res 75:172–181. doi: 10.1002/jnr.10857

55. Vertes RP (2004) Differential Projections of the Infralimbic and Prelimbic Cortex in the Rat. Synapse 51:32–58. doi: 10.1002/syn.10279

56. Vincent S, Khan Y, Benes F (1993) Cellular distribution of dopamine D1 and D2 receptors in rat medial prefrontal cortex. J Neurosci 13:2551–2564. doi: 10.1523/JNEUROSCI.13-06-02551.1993

57. Zhou T, Sandi C, Hu H (2018) Advances in understanding neural mechanisms of social dominance. Curr Opin Neurobiol 49:99–107. doi: 10.1016/j.conb.2018.01.006

58. Zhou T, Zhu H, Fan Z, Wang F, Chen Y, Liang H, Yang Z, Zhang L, Lin L, Zhan Y, Wang Z, Hu H (2017) History of winning remodels thalamo-PFC circuit to reinforce social dominance. Science 357:162–168. doi: 10.1126/science.aak9726

